# Kidney medulla macrophages maintain a free flow of urine by sensing force

**DOI:** 10.64898/2026.07.02.736225

**Authors:** Rukun He, Zhanghui Huang, Yanran Li, Jian He, Guo Cheng, Qiang Wang, Ningting Chen, Yuancheng Weng, Xinge Wang, Xiaoli Liu, Xiao Z Shen

## Abstract

Blockade by sedimentary particles, such as mineral crystals, is a continuous risk the kidney tubule faces. To prevent that, kidney resident macrophages form transepithelial protrusions and remove intratubular sedimentary particles, a behavior particularly prevailing in the medulla over the cortex. However, the molecular mechanisms underlying this characteristic behavior of medulla macrophages are incompletely understood. In this study, we identified that the medulla had higher mechanical stiffness than the cortex in steady state, which was further elevated when kidney stone formed. Increased tissue rigidity was sensed by medulla macrophages via mechanoreceptor Piezo1, which promoted macrophage protrusion formation and their ability to clean the tubules. Loss of Piezo1 expression in kidney macrophages predisposed mice to intratubular accumulation of mineral crystal in steady state and accelerated kidney stone formation during oxalate intake challenge. Signaling via Piezo1 mobilized molecules involved in cell adhesion and protrusion assembly, including Talin2 and focal adhesion kinase (FAK). Finally, we developed a first-of-its-kind cell-based therapy for the treatment of experimental nephrolithiasis by exploiting macrophage Piezo1 activity, and this strategy shows great promise for future translational research.

## INTRODUCTION

The kidney tubule system has one of the most mechanically diverse environments in the body. On one hand, kidney tubules drain glomerular filtrates, the amount (speed) of which is determined by renal blood flow, plasma colloid osmotic pressure and Bowman’s capsule hydrostatic pressure (1). On the other hand, with water being reabsorbed when glomerular filtrates passing through the tubule system, various particles including mineral crystals are generated due to supersaturation of urine solutes (2). Overgeneration of sedimentary particles would partially or completely block tubules, leading to tubule dilation and mechanic injury. Mineral crystals would aggregate and grow, gradually turning into kidney stones. Thus, tubule occlusion by sedimentary particles is the most imminent stress the kidney faces and up to 1/4 of fresh urine samples from healthy subjects show overt crystalluria (3). However, Pharmacological treatment options for nephrolithiasis remain limited, and renal stone removal predominantly relies on surgical interventions such as extracorporeal shock wave lithotripsy (ESWL) (4).

Thus, routine removal of intratubular sedimentary particles is critical to assure free urine flow and normal kidney functions. However, urine flush is not enough to keep tubule unobstructed, and additional cellular mechanism is required. It was recently shown that kidney medulla macrophages (MØ) constitutively formed transepithelial protrusions and facilitated particle removal from the urine (5). Depleting MØ or blockade their integrin interaction with the tubules would significantly accelerate kidney stone formation (5). This finding provides an unprecedented opportunity for developing cell therapeutics for nephrolith diseases.

Mineral crystallization due to urine supersaturation is particularly concerning tubules in the downstream medulla over the upstream cortex. The formation of transepithelial protrusions is more pronounced in the medulla MØ relative to cortex MØ (5), suggesting a medulla-specific environmental cue driving local resident MØ to fulfill tissue-specific needs and handle tissue-specific challenges. One of the microenvironmental cues is cell–extracellular matrix (ECM) interaction which provides structural, biochemical, and signaling support, regulating the intracellular cytoskeletal organization, cell polarity and functions (6). In the current study, we further explored the mechanism underlying the formation of transepithelial protrusions by medulla MØ and identified that kidney medulla has high stiffness relative to cortex. Medulla MØ but not cortex MØ highly expressed the mechanosensitive cation channel Piezo1 which drove transepithelial protrusion formation by medulla MØ. Loss of Piezo1 in medulla MØ predisposed mice to kidney stone formation, while delivery of a Piezo1 activator could reduce kidney stone formation in both prophylactic and curative approaches in the setting of hyperoxaluria insult.

## RESULTS

### Medulla macrophages highly express the mechanosensor Piezo1

Previous work showed that tubular obstruction by kidney stone formation resulted in tubule dilation, which was accompanied with an enhanced formation of transepithelial protrusions by medulla MØ (5), implying that mechanosensing may prompt the generation of transepithelial protrusions by medulla MØ. To interrogate this hypothesis, we first evaluated and compared the expression profiles of mechanoreceptors between kidney cortex and medulla MØ since morphological difference between them already existed in steady state, as medulla MØ generally had greater protrusion complexity (Figure 1A) (5). Transcript analysis revealed that medulla MØ highly expressed *Piezo1* and *Trpv4* but had limited expression of other mechanoreceptors (Figure 1B). This selective expression pattern was confirmed by our transcriptomic study of medulla MØ (Figure 1C). Interestingly, only *Piezo1* expression displayed a remarkable difference between medulla and cortex MØ favoring the former (Figure 1B). Further considering that Piezo1 detects external mechanical forces while *Trpv4* encodes an osmotic- and temperature-gated ion channel, we next focused on the expression and function of Piezo1 in medulla MØ. We first confirmed that medulla MØ highly expressed Piezo1 over cortex MØ in protein level by flow cytometry (Figure 1D). Functionally, we verified that Piezo1-selective activator Yoda1 could induce Ca^2+^ influx in medulla MØ, manifested by recording medulla MØ derived from *CX3CR1*^CreERT2/+^: GCaMP6f mice ex vivo (Figure 1E and Video S1). In addition, mechanical brushing caused a transient calcium wave in the medulla MØ within the first 30 seconds, which was completely abrogated in Piezo1-deprived MØ derived from *CX3CR1*^CreERT2/+^: GCaMP6f: *Piezo1*^fl/fl^ mice (Figure 1F), confirming that Piezo1 expressed on kidney MØ mediated pressure-induced alterations. Together, these results substantiated that Piezo1 expressed by medulla MØ could convert physical forces into biochemical signals.

**Figure 1.**
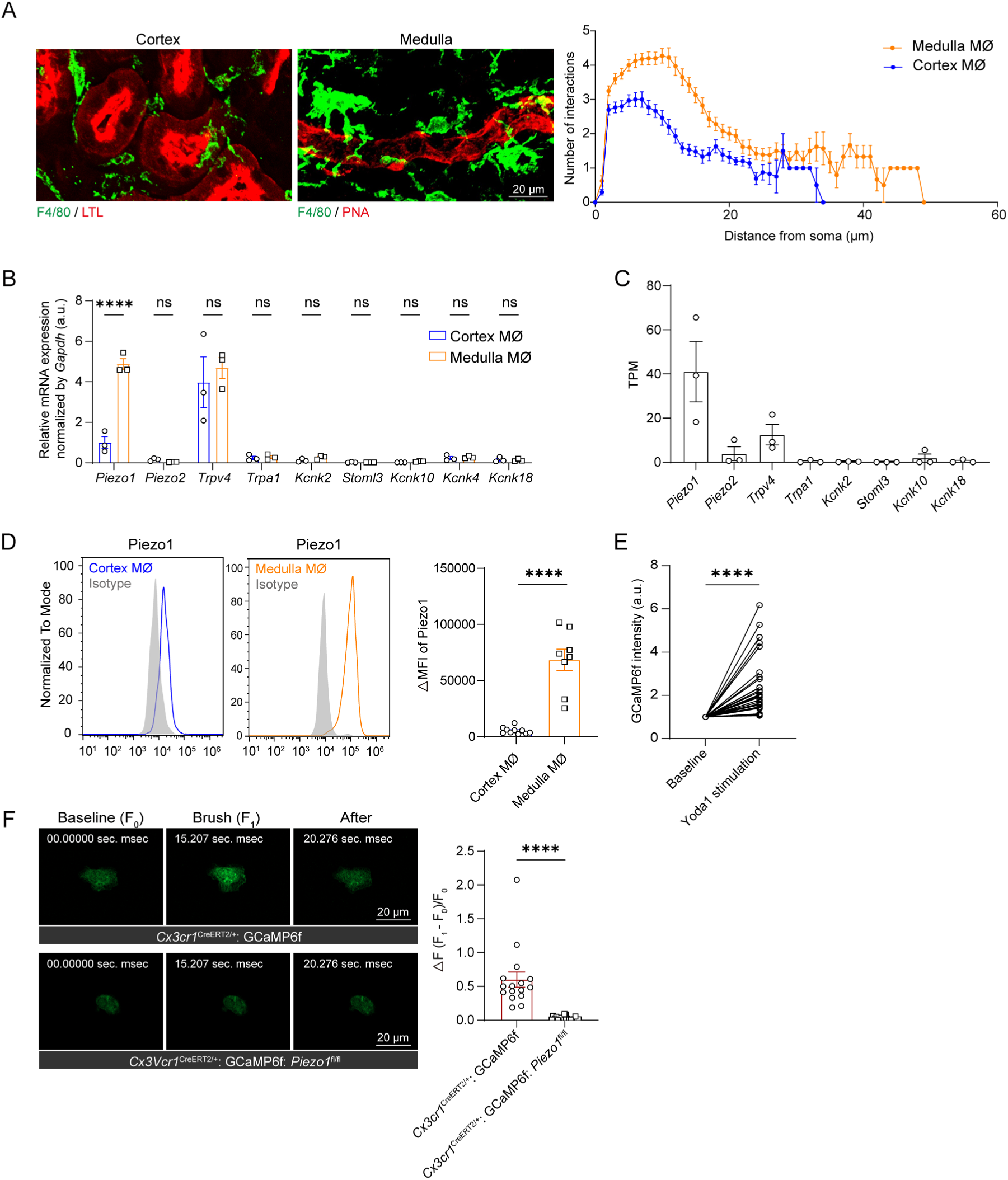
Medulla macrophages highly express Piezo1. (**A**) Confocal images showing morphological differences of F4/80^+^ MØ between the cortex and medulla (left). Quantification of MØ protrusion complexity by Sholl analysis (right). n = 6. (**B**) The transcript expression profiles of mechanoreceptors in cortex and medulla MØ, assessed by RT-PCR. Each dot represents a pool of cells derived from 4 mice. (**C**) The relative expression levels of all known mechanoreceptor genes in medulla MØ derived from *CX3CR1*^CreERT2/+^ control mice of the transcriptomic study described in Figure 5A, according to gene expression values (TPM) of bulk RNA-seq. (**D**) Piezo1 expression in cortex and medulla MØ, evaluated by flow cytometry. (**E**) Relative GCaMP6f fluorescence intensity in medulla MØ purified from tamoxifen-treated *CX3CR1*^CreERT2/+^: GCaMP6f mice before and after Yoda1 stimulation. Each dot represents an individual cell. (**F**) Representative time-lapse frames taken from GCaMP6f fluorescence recordings of medulla MØ from *CX3CR1*^CreERT2/+^: GCaMP6f mice and *CX3CR1*^CreERT2/+^: GCaMP6f: *Piezo1*^fl/fl^ mice before and after mechanical brushing. Quantification of peak amplitude (ΔF/F) of individual medulla MØ in response to brushing. Each dot represents an individual cell. n = 4. n.s., not significant. \*\*\*\**P* < 0.0001 by two-way ANOVA in (B) and two-tailed unpaired t test in (D, F), and two-tailed paired t test in (E). Data are depicted as mean ± SEM. Data are derived from at least 2 independent experiments.

### Kidney medulla has a stiffer microenvironment than kidney cortex

Increasing substratum stiffness would upregulate Piezo1 expression (7–9). Indeed, we noticed a further upregulation of Piezo1 expression by medulla MØ in an oxalate nephrolith model which induced kidney stone formation preferentially in the medulla (Figures 2A and S1).

**Figure 2.**
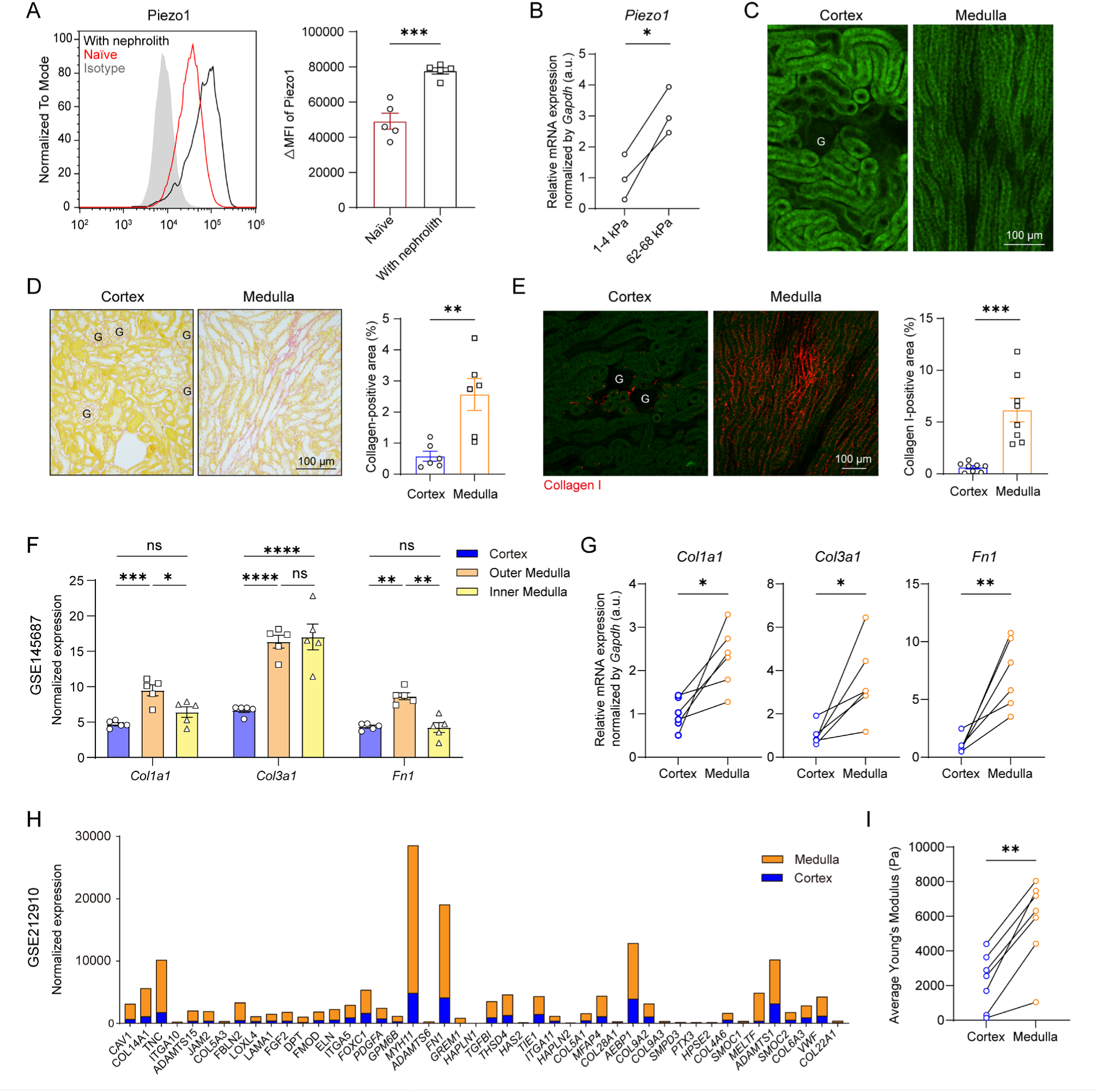
Kidney medulla has higher rigidity relative to cortex. (**A**) Oxalate nephrolith model was induced by treating mice i.p. with NaOx (60 mg/kg), as shown in Supplementary figure 1. Piezo1 expression change in medulla MØ between naïve mice and mice with nephrolith was evaluated on day after by flow cytometry. (**B**) Cortex MØ were purified and then plated on hydrogels with different stiffness. After 24 h, cells were subject to RT-PCR analysis for evaluating *Piezo1* expression. Each dot represents a pool of cells derived from 4 mice. We did not examine Piezo1 expression by flow cytometry because trypsin digestion for cell detachment impaired Piezo1 detectability. (**C**) The morphology of tubules in the cortex and medulla. Tubules are readily identifiable by their autofluorescence. (**D**) Picrosirius red staining showed collagen fiber deposition in the cortex and medulla. Each dot represents the average of two 696 × 524 μm^2^ fields of view (FOVs) from one mouse. (**E**) Immunohistostaining showed collagen I in the cortex and medulla. Each dot represents the average of two 649 × 649 μm^2^ FOVs from one mouse. (**F**) Normalized gene expression of *Col1a1*, *Col3a1*, and *Fn1* in the cortex and different positions of medulla from a GEO dataset of mice (GSE145687). (**G**) Transcript analysis of *Col1a1*, *Col3a1* and *Fn1* in the cortex and medulla by RT-PCR. (**H**) Normalized gene expression of all genes included in the GO term “Extracellular matrix organization” (GO: 0030198) in human kidney cortex and medulla, derived from a GEO dataset (GSE212910). (**I**) Average Young’s modulus (Pa) of cortex and medulla measured by atomic force microscopy. n.s., not significant. \**P* < 0.05, ** *P* < 0.01, *** *P* < 0.001, **** *P* < 0.0001 by two-tailed unpaired t test (A, D, E), two-tailed paired t test (B, G, I) and two-way ANOVA (F). Data are depicted as mean ± SEM. Data are derived from at least 2 independent experiments.

Interestingly, cortex MØ would also upregulate *Piezo1* expression when the matrix rigidity increased in an in vitro culture system (Figure 2B), suggesting that the difference between medulla and cortex MØ in Piezo1 expression may result from discrepancy in tissues stiffness. We were thus set to investigate the difference of tissue stiffness between kidney cortex and medulla by multiple approaches. By morphology, tubules in the medulla were generally straight and rigid while those in the cortex were convoluted, implying a relatively higher degree of stiffness in medulla (Figure 2C). It is well accepted that disposition of the extracellular matrix (ECM) determines the stiffness of tissue. In our study, picrosirius red staining showed that more collagen fibers distributed in the ECM of medulla over cortex in the kidneys of normal C57BL/6 mice (Figure 2D). This was consistent with our result of immunohistostaining of collagen I (Figure 2E). Moreover, mining the transcriptomic dataset from a mouse kidney study (10) revealed that fibrotic genes *Col1a1*, *Col3a1* and *Fn1* were over-represented in the medulla relative to cortex (Figure 2F), which was in line with our RT-PCR study (Figure 2G). These findings were cross-validated by searching published transcriptomic datasets of human (11) which showed that ECM gene products were generally enriched in the medulla over cortex (Figure 2H). To directly measure the stiffness of kidney tissue, we performed atomic force microscopy indentation experiments on mouse kidney slices. It showed that kidney medulla was ∼2.6-fold higher than cortex in stiffness (Figure 2I). Altogether, these data indicated that kidney medulla has a more rigid microenvironment relative to the kidney cortex, correlating with a higher expression level of Piezo1 in medulla MØ over their cortex counterparts.

### Piezo1 is required by medulla macrophages for preventing kidney stone formation

To interrogate the significance of MØ Piezo1 in the formation of transepithelial protrusions and its related role in maintaining tubule homeostasis, we crossed *Cx3cr1*^CreERT2/+^ mice with *Piezo1*^fl/fl^ mice (Figure S2A). Intraperitoneal tamoxifen treatment reduced *Piezo1* transcripts by ∼70% in medulla MØ of the offspring (hereafter *Piezo1*^ΔMØ^) (Figure S2C). Before that, *Cx3cr1*^GFP/+^ reporter mice had verified that CX3CR1 highly expressed in MØ but not in medullary tubules (Figure S2B) (5) Interestingly, acute deprivation of Piezo1 expression in medulla MØ markedly altered the morphology of medulla MØ, with shortened protrusions and less protrusion complexity, in comparison to tamoxifen-treated control *Cx3cr1*^CreERT2/+^ mice (Figures 3A-C). In addition, a significant reduction in the density of transepithelial protrusions was noticed, and actually most medulla MØ appeared detached from renal tubules (Figures 3A and 3D). Transepithelial protrusions were defined by their penetration through the Peanut Lectin Agglutinin (PNA)^+^ epithelial cells in confocal analysis of thicker sections (30 μm) of the medulla with high-resolution and optical magnification (Figure E), as shown before (5). To investigate whether stimulating Piezo1 had a direct effect on MØ in protrusion formation, we purified medulla MØ and stimulated them with Yoda1 ex vivo. Coherent with the in vivo finding, Yoda1-pretreated medulla MØ derived from reporter *Cx3cr1*^GFP/+^ mice displayed an accelerated protrusion extension upon being re-plated on fibronectin substratum, when compared to vehicle-pretreated MØ which remained generally spherical in the 1^st^ hr (Figure 3F). We used fibronectin-coated substratum because integrin β1-fibronectin ligation is required by medulla MØ to form transepithelial protrusions (5). These data together suggest that mechanosensing by Piezo1 relays a high environmental stiffness cue to facilitate a superior protrusion growth in kidney medulla MØ.

**Figure 3.**
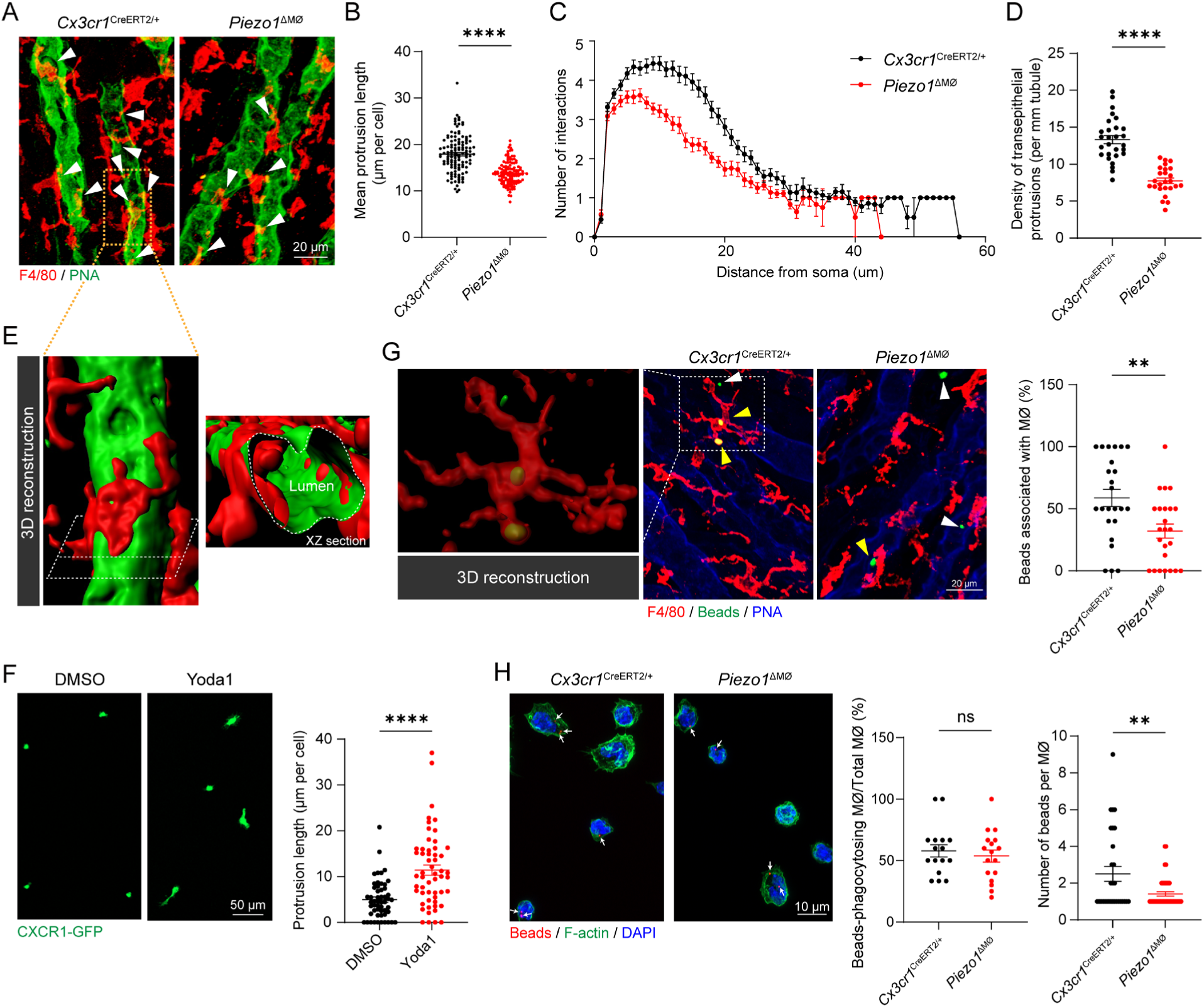
Loss of Piezo1 impairs protrusion formation and bead-sequestering ability of medulla macrophages. (**A-D**) The morphology of medulla MØ and their relationship with collecting ducts in *Piezo1*^ΔMØ^ mice and the control *Cx3cr1*^CreERT2/+^ mice. (A) Representative images showing MØ (red) and their relationship with the medullary collecting ducts (green). Arrowhead, transepithelial protrusion. Z-projections of 20 μm. (B) Length of protrusion, (C) complexity of protrusion, and (D) density of transepithelial protrusions of MØ were compared between these two groups. n = 6. Each dot represents an individual cell in (B). Each dot represents the quantification of one 319 × 319 μm^2^ FOV in (D). (**E**) Confocal image of the spatial relationship between PNA^+^ epithelium and F4/80^+^ cell protrusions. The magnifications show the corresponding 3D reconstructions viewed from abluminal (left) and luminal (right) sides. Z-projections of 20 μm. (**F**) Medulla MØ purified from *Cx3cr1*^GFP/+^ mice were pretreated with Yoda1 or vehicle for 24 hr. Cells were then replated on fibronectin-coated plate for 1 hr and the length of newly grown protrusions was measured and compared. n = 6. (**G**) Mice received intrapelvic injection of fluorescent latex beads. After 8 h, the interactions between MØ (red) and beads (green) were examined around medullary collecting ducts (blue). Z-projections of 20 μm. Each dot represents the quantification from one 319 × 319 μm^2^ FOV. n = 6. (**H**) Medulla MØ were purified from the indicated mice and were then co-incubated with latex beads (red) for 24 h. The frequency of bead-containing MØ and the number of beads in bead-containing MØ were enumerated. F-actin (green) and DAPI (blue) staining displayed the area of cytoplasm and nucleus of MØ, respectively. Each dot represents the quantification from one FOV. n = 4. n.s., not significant. \*\**P* < 0.01; \*\*\*\**P* < 0.0001 by two-tailed unpaired t test. Data are depicted as mean ± SEM. Data are derived from at least 2 independent experiments.

Considering that medulla MØ actively facilitate in removing intratubular sedimentary particles with the help of their transepithelial protrusions (5), we posited that the reduced degree of protrusion formation when MØ Piezo1 expression was abrogated would increase the deposition of sedimentary particles in renal tubules. To test this hypothesis, we first injected fluorescent inert latex beads (0.5 μm in diameter) into kidney pelvis. Some beads would be backflushed into the upstream collecting ducts of kidneys and previously we showed that medulla MØ could phagocytosed intratubular beads (5). At 8 h post infusion, the quantities of beads in the medulla were comparable between *Piezo1*^ΔMØ^ and the control mice (Figure S3). While over 50% of intrarenal beads were associated with MØ, including those phagocytosed by MØ, in the control mice, most of the beads in the *Piezo1*^ΔMØ^ mice were intraluminal in the tubules and fewer beads were associated with medulla MØ (Figure 3G). To test whether loss of Piezo1 also impaired MØ phagocytosis, we purified medulla MØ and fed them with fluorescent beads for 24 hr in vitro. It displayed that although the frequency of bead-phagocytosed cells was comparable between the two genotypes, *Piezo1*-deprived MØ had reduced ability in taking up multiple beads when compared to *Piezo1*-intact MØ (Figure 3H). Taken together, these data suggest an impaired tubule cleaning function of medulla MØ in the absence of Piezo1 expression.

The urine typically contains a significant number of aggregated particles derived from varying sources. The most pronounced source is mineral microcrystals driven by supersaturation, which occurs as substantial water reabsorption takes place while glomerular filtrate flows through the tubule system. Calcium phosphate (CaPi) is one of the major mineral crystal types (12). Surprisingly, deprivation of MØ Piezo1 expression (Figure 4A) resulted in a widespread deposition of CaPi microcrystals in medulla tubules, which was absent in the control mice. This was revealed by both OsteoSense (a fluorescent bisphosphonate that can specifically bind to crystalline CaPi) (13) (Figure 4B) and Von Kossa staining (for calcium crystals) (Figure S4A). To be noted, in *Piezo1*^ΔMØ^ mice, the concentrations of Ca^2+^ or Pi in blood plasma was not altered (Figure S4B); neither urine output nor the excretion of Ca^2+^ or Pi was changed (Figure S4C). In addition, the expression of *Slc34a1*, the major phosphate transporters responsible for reabsorbing Pi from the urine, in proximal tubules, was intact (Figure S4D).

**Figure 4.**
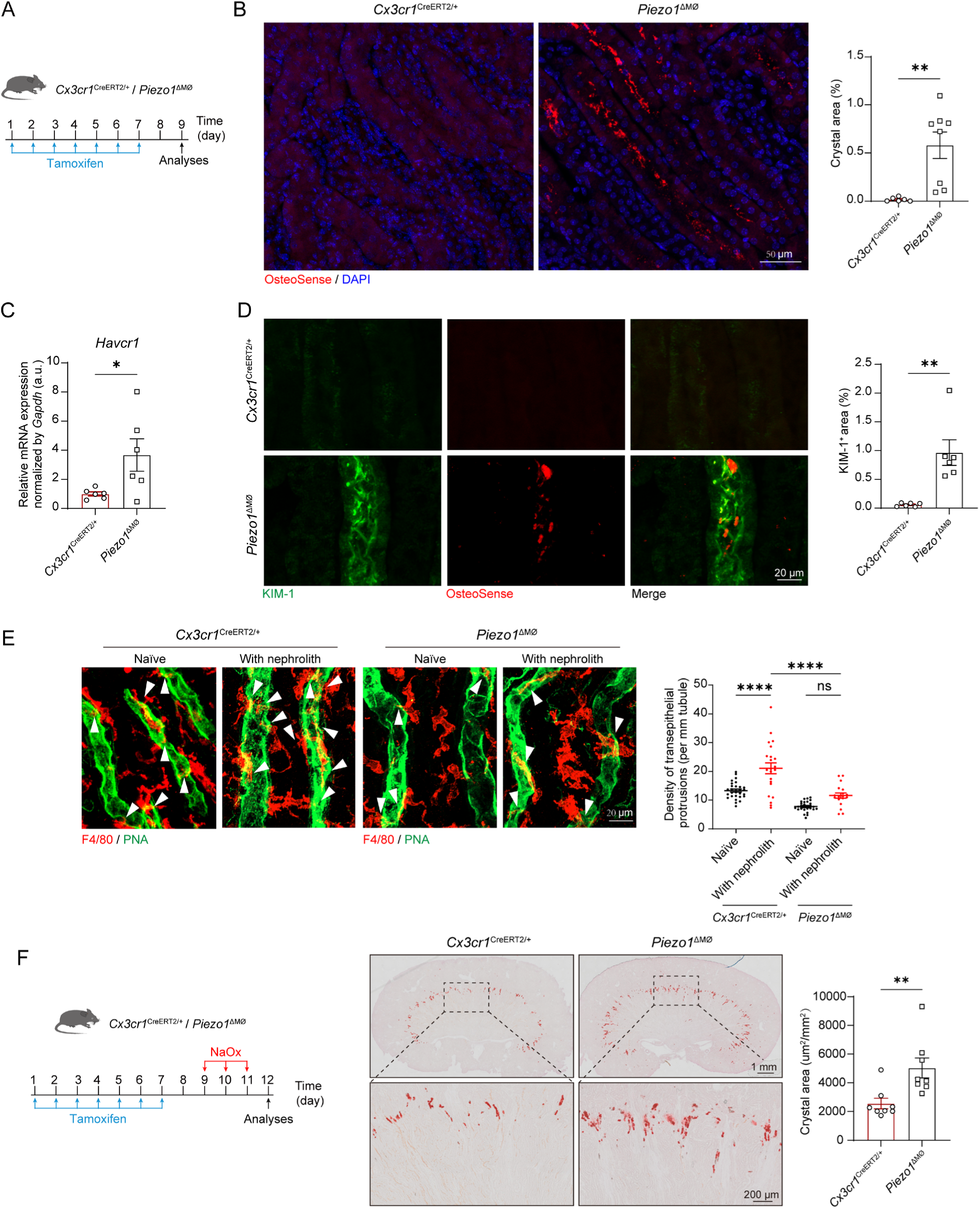
Loss of Piezo1 in medulla macrophages predisposes mice to mineral stone formation. (**A**) Scheme illustrating the protocol of experiments for Panels B-D. (**B**) CaPi particles (red dots) highlighted by OsteoSense detection. CaPi crystal areas in the medulla were measured and compared. Each dot represents the average of the affected areas of two 319 × 319 μm^2^ FOVs from one mouse. (**C**) *Havcr1* expression in kidneys medulla were evaluated by RT-PCR analysis. (**D**) KIM-1 expression in the tubules around CaPi crystals (OsteoSense^+^) of *Piezo1*^ΔMØ^ mice. Each dot represents the average of the affected areas of two 319 × 319 μm^2^ FOVs from one mouse. (**E**) With an acute hyperoxaluria model, confocal images of MØ (red) and the collecting ducts (green) in the medulla at hour 4 post the NaOx (60 mg/kg) treatment, with Z-projections of 20 μm. Quantification of the densities of the transepithelial protrusions (arrowheads) was shown. Each dot represents the enumeration of one 319 × 319 μm^2^ FOV. n = 6. (**F**) Scheme illustrating the protocol of oxalate nephrolith experiments (left). Kidney stone deposition was examined by Alizarin Red S (right). Quantification of the affected area in the whole kidney was also shown. Each dot represents the average of the affected areas of two whole-mount slides from one mouse. n.s., not significant. \**P* < 0.05, ** *P* < 0.01; **** *P* < 0.0001 by two-tailed unpaired t test in (B, C, D, F) and two-way ANOVA in (E). Data are depicted as mean ± SEM. Data are derived from at least 2 independent experiments.

Mineral crystals have been shown to injure tubular epithelial cells (14, 15). The increased intratubular crystal deposition in *Piezo1*^ΔMØ^ mice was indeed accompanied with an increased expression of renal epithelial injury markers *Havcr1* which encodes KIM-1, as assessed by RT-PCR (Figure 4C). The conclusion that tubule injury resulted from crystal deposition was further highlighted by our observation that KIM-1^+^ injured tubular epithelial cells were those surrounding CaPi crystals (Figure 4D). These data underscored tubule clearance by medulla MØ is critical in preventing collateral damages to tubules due to mineral crystal deposition.

CaOx is the most prevalent type of kidney stone in human. To gain insight into the impact of MØ on CaOx stone development, we first induced an acute CaOx nephrolith model by giving one i.p. shot of NaOx (5). As nephroliths increase the stiffness of kidney medulla, an increase of MØ-derived transepithelial protrusions in the medullary collecting ducts was noticed in the control mice (Figure 4E). However, the trend of increased transepithelial protrusions upon CaOx crystal formation was significantly suppressed in *Piezo1*^ΔMØ^ mice (Figure 4E). Consequently, deprivation of MØ Piezo1 during a continuous hyperoxaluria challenge accelerated kidney stone formation (Figure 4F). Altogether, there data underscore an indispensable role of Piezo1 in mediating the function of medulla MØ in removing intratubular sedimentary particles and in maintaining the integrity of kidney tubules.

### Piezo1 activation promotes protrusion formation

To interrogate the roles of Piezo1 in medulla MØ biology, we next took a transcriptomic approach by performing bulk RNA-seq analysis of medulla MØ derived from adult *Piezo1*^ΔMØ^ mice and the control *Cx3cr1*^CreERT2/+^ mice after tamoxifen treatment (gating strategy shown in Figure S5A (5, 16). Hierarchical clustering analysis showed that the replicates of each cell subset were clustered together (Figure S5B). A distinction in the transcriptomes between medulla MØ of disparate *Piezo1* expression was unveiled, as 1,363 genes were differentially expressed (fold change > 2; p value < 0.05) (Figure 5A). Among the differentially expressed genes (DEGs), downregulated genes overwhelmed upregulated genes (1,083 vs. 280) in the *Piezo*1-deprived cells. Gene ontology (GO) enrichment for biological process showed that the *Piezo*1-deficient medullary MØ were under-represented with genes associated with membrane potential (Figure 5B), which was consistent with the identity of Piezo1 as a mechanically activated ion channel. Interestingly, GO analysis highlighted functional impairment in cell adhesion, synapse assembly and cell junction assembly by *Piezo*1-deficient MØ (Figure 5B), which was consistent with their phenotype of detachment from the tubules and weakened transepithelial protrusion formation (Figures 3A and 3D). Piezo1 activation was shown to promote efferocytosis (17, 18) and inflammation (19–22) by MØ of non-renal origin. However, phagocytosis- or inflammation-related terms did not appear in the top 20 downregulated GO terms of MØ deprived of *Piezo1*, implying MØ tissue-specificity. Genes involved in smooth muscle proliferation was over-represented in the Piezo1-deficient medulla MØ relative to their controls (Figure S5C), which appeared irrelevant to our current study.

**Figure 5.**
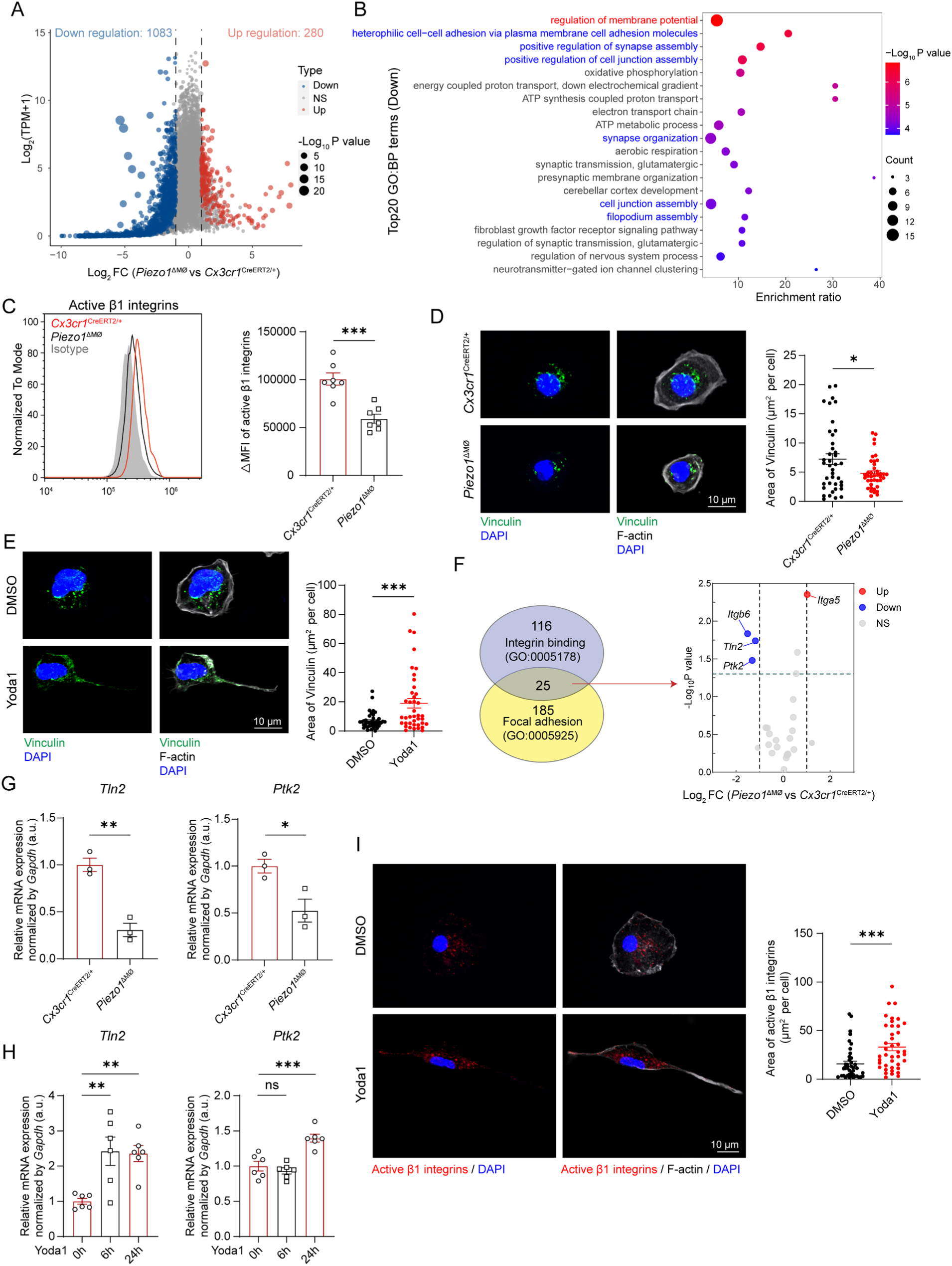
Piezo1 activation mobilizes the machinery for protrusion formation. (**A**) Volcano plot shows statistical significance when comparing kidney medulla MØ in the absence of Piezo1 vs. in the presence of Piezo1. Genes upregulated in *Piezo1-deprived* MØ are represented by red symbols, whereas those downregulated are shown in blue. (**B**) Top 20 downregulated pathways of *Piezo1-deprived* medulla MØ relative to control cells, derived from GO analysis of DEGs. The term about membrane potential is shown in red, and the terms about cytoskeleton mobilization are highlighted in blue. (**C**) Medulla MØ from *Piezo1*^ΔMØ^ mice and the control mice were subject to flow cytometry analysis of activated integrin β1. (**D**) Medulla MØ were purified from *Piezo1*^ΔMØ^ mice and their control mice. After being plated on fibronectin-coated plate for 2 hr, focal adhesion density per cell, as assessed by Vinculin staining, was shown. F-actin was co-stained to display the general cell morphology. Each dot represents an individual cell. n = 6. (**E**) Medulla MØ purified from C57/BL6 mice were pretreated with Yoda1 or vehicle for 24 hr. Cells were then replated on fibronectin-coated plate for 2 hr. Focal adhesion density was then evaluated. Each dot represents an individual cell. n = 6. (**F**) From the 116 and 185 genes respectively belonging to GO terms “integrin binding” and “focal adhesion”, there were 25 genes overlapping. Among them, the upregulated and downregulated genes in the *Piezo1*-deficient MØ relative to control MØ were shown. The genes with fold change > 2 and p value < 0.05 were highlighted. (**G**) *Tln2* and *Ptk2* expression in medulla MØ of the indicated mice were evaluated by RT-PCR. Each dot represents a pool of cells derived from 4 mice. (**H**) BMDM were activated with Yoda1 for the indicated time, and *Tln2* and *Ptk2* expression were evaluated by RT-PCR. (**I**) BMDM were activated with Yoda1 for 24 h, and the activated integrin β1 was evaluated by immunocytofluorescent staining using a specific anti-integrin β1 antibody (Clone 9EG7). We did not use flow cytometry to measure because the conformation of integrin β1 would be altered after digestion by trypsin and EDTA during cell detachment. Each dot represents an individual cell. n = 4. n.s., not significant. * *P* < 0.05, ** *P* < 0.01; *** *P* < 0.001 by two-tailed unpaired t test in (C, D, E, G, I) and one-way ANOVA in (H). Data are depicted as mean ± SEM. Data are derived from at least 2 independent experiments.

Previously we showed that interaction between MØ integrin β1 with fibronectin at basolateral side of tubules was required for the formation of transepithelial protrusions (5). Piezo1 did not appear to promote MØ integrin β1 expression since neither our RNA-seq data nor RT-PCR analysis of medulla MØ indicated an alteration of *Itgb1* expression after *Piezo1*-abrogation (Figures S5D and S5E). In addition, integrin β1 expression at protein level did not change, as manifested by flow cytometry analysis using an antibody recognizing total integrin β1 (Figures S5F). Conformation shift from inactivated state to activated state is integral to integrin β1 functions (23). Employing an antibody specific to activated integrin β1 (Clone 9EG7) (24, 25), we detected a diminished activity of integrin β1 on *Piezo*1-deficient MØ (Figure 5C), suggesting that *Piezo1* abrogation disrupted inside-out signaling for activating integrin(s) (26, 27). Integrin-based focal adhesion to extracellular matrix is important for protrusion formation (6, 28). Congruent with this notion, Piezo1-deficient medulla MØ exhibited decreased focal adhesion formation ex vivo, as assessed by Vinculin staining, when they were plated in fibronectin-coated plate for 2 h (a time point when MØ had firmly attached to the plate) (Figure 5D). In contrast, activating Piezo1 with Yoda1 promoted focal adhesion formation in medulla MØ derived from wild-type mice (Figure 5E). These data together could at least partially explain the detachment of medulla MØ from tubules in *Piezo1*^ΔMØ^ mice, which was in alignment of a decreased density of transepithelial protrusions (Figures 3A and 3D).

To further explore the molecular events underlying Piezo1-facilitated protrusion formation in medulla MØ, we analyzed our RNA-seq data, particularly focusing on the genes associated with GO terms “integrin binding” and “focal adhesion”. There were 25 genes included in both terms of Gene Ontology database. Among them, *Itgb6*, *Tln2* and *Ptk2* genes were distinguishably downregulated in the *Piezo1*-deficient MØ (Figure 5F). Considering that *Itgb6* expression was relatively low in comparison to most other integrin β subtypes (Figures S5G), we did not investigate *Itgb6* further. *Tln2* and *Ptk2* genes encode Talin2 and focal adhesion kinase (FAK), respectively. Talins are essential for integrin activation and signaling, which triggers a cascade of internal signals, most notably through FAK which serves as a signaling hub (29–32). We verified that both *Tln2* and *Ptk2* transcripts were downregulated in *Piezo1*-deficient MØ (Figure 5G). Of note, the expression of *Tln1* which encodes Talin1 was not altered in MØ deprived of *Piezo1* expression (Figures S5H). Bone marrow-derived MØ (BMDM) from C57BL/6 mice highly express Piezo1 as well (Figure S6). Using BMDM as surrogate cells to kidney medulla MØ, we showed that Yoda1 stimulation could induce wild-type MØ to upregulate *Tln2* and *Ptk2* expression (Figure 5H), which was accompanied with more integrin β1 in an activated state (Figure 5I).

To be noted, the response of *Tln2* upregulation by MØ upon Yoda1 stimulation was faster and more pronounced than that of *Ptk2* upregulation, we thus focused on Talin2 for further investigation. Stimulation of Piezo1 results in Ca^2+^ influx. Consistently, blocking calpain, the downstream player relaying Ca^2+^ signal, could deprive Yoda1’s effect on *Tln2* expression in BMDM (Figure 6A). Flow cytometry analysis verified that more Talin2 expression in medulla MØ over their cortex counterparts in wild-type kidney (Figure 6B). In alignment, Talin2 was downregulated in *Piezo1*-deprived medulla MØ relative to controls (Figure 6C). In contrast, kidney stone formation which increased tissue stiffness was accompanied with an increase of Talin2 expression in medulla MØ of wild-type mice (Figure 6D). Immunocytofluorescent analysis of wild-type medulla MØ showed that in normal state, Talin2 generally did not overlap well with F-actin, as it was centralized around the nucleus while F-actin relatively aggregated at periphery (Figure 6E). Piezo1 activation by Yoda1 promoted cell polarity, accompanied with not only an increased Talin2 expression but also an altered distribution of Talin2, as it started to distribute along the extended protrusions together with F-actin (Figure 6E). Focal adhesions consist of peripheral small form and centrally located large form (6, 33, 34). In the cells without Yoda1 treatment, Talin2 appeared partially colocalized with the centrally located large focal adhesions but has no distribution in the area of peripheral small focal adhesions (Figure 6F). After Piezo1 activation by Yoda1, the central focal adhesions turned even larger and Talin2 accumulated in their vicinity (Figure 6F). Moreover, some Talin2 was then at least partially aligned with peripheral focal adhesions into the extended protrusions (Figure 6F). These data together suggest a mobilized cell skeleton promoting protrusion formation in response to Piezo1 stimulation.

**Figure 6.**
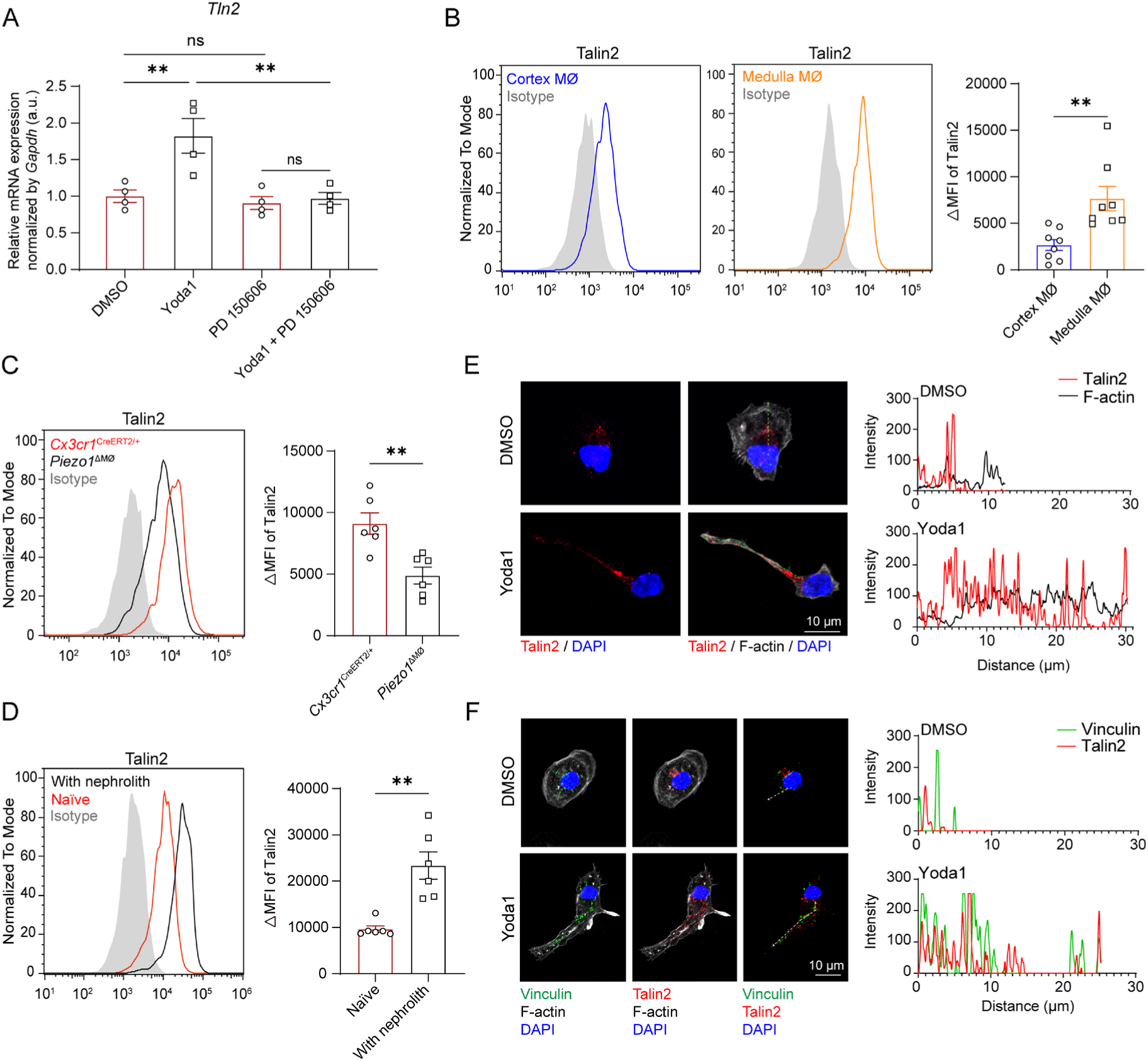
Piezo1 activation promotes Talin2 expression and its distribution along protrusion. (**A**) BMDM were stimulated with Yoda1 for 24 h, in the presence or absence of PD 150606, a calpain inhibitor. Some cells were treated with DMSO, the vehicle of Yoda1. *Tln2* expression was evaluated by RT-PCR. (**B**) MØ purified from kidney cortex and medulla were evaluated for Talin2 expression by flow cytometry. (**C**) MØ purified from the medulla of *Piezo1*^ΔMØ^ mice and their control mice were evaluated for Talin2 expression by flow cytometry. (**D**) C57BL/6 mice were i.p. treated with NaOx (60 mg/kg) to develop kidney stones. One day later, kidney medulla MØ were purified and evaluated by flow cytometry for Talin2 expression, in comparison to the counterparts from naïve mice. (**E** and **F**) Medulla MØ purified from C57/BL6 mice were pretreated with Yoda1 or vehicle for 24 hr. Cells were then replated on fibronectin-coated plate for 2 hr. (**E**) Representative images show the distribution of Talin2 (red) and F-actin (white) under DMSO and Yoda1 treatment. Pearson’s correlation coefficients were 0.06 ± 0.07 for the DMSO group and 0.22 ± 0.04 for the Yoda1 group. Intensity traces (offset green dotted line which indicates the longest distance from nucleus to cell membrane) are plotted at right. Pearson’s correlation coefficient represent means ± SEM from 24–30 cells per group. n = 6. (**F**) Representative images show the distribution of Vinculin (green) and Talin2 (red) under DMSO and Yoda1 treatment. Pearson’s correlation coefficients were 0.31 ± 0.08 for the DMSO group and 0.61 ± 0.04 for the Yoda1 group. Intensity traces (offset white lines which indicates the longest distance from nucleus to cell membrane) are plotted at right. Pearson’s correlation coefficient represent means ± SEM from 24–30 cells per group. n = 6. n.s., not significant. \*\**P* < 0.01 by one-way ANOVA in (A) and two-tailed unpaired t test in (B, C, D). Data are depicted as mean ± SEM. Data are derived from at least 2 independent experiments.

### Employing kidney macrophages to develop cell therapy for nephrolithiasis in mice

Nephrolithiasis has very limited pharmacological options and there is an unmet need for non-invasive therapeutics. It is therefore tempting to translate our findings about Piezo1-mediated regulation of medulla MØ into a valid therapeutic strategy, i.e., a cell-based therapy, for nephrolithiasis. We recently reported that a retrograde delivery via non-invasive intravesical route offered a promising strategy for targeting kidney, which had limited off-target effects on tissues/organs outside of the urinary system (35). We postulated that via this route, a Piezo1 agonist could be delivered to the kidney tubule system, which would favor the formation of transepithelial protrusion by medulla MØ and had minimal side effects on extrarenal organs. Indeed, after intravesical delivery of Yoda1 (a 355 Da small molecule) to female mice, we could detect Yoda1 in the kidney at 8 h post intravesical treatment (the time point at which peak concentration reached in the kidney post intravesical delivery, referring to the findings shown in Reference (35)) by mass spectrometry, while Yoda1 was not detectable in the heart (Figures S7A). Moreover, leakage of Yoda1 to the blood was minor compared to any given time after an intraperitoneal delivery (Figures S7B). Thus, intravesical infusion of Yoda1 is kidney specific with little leakage to extrarenal organs/tissues.

Intravesical delivery of Yoda1 to wild-type mice once a day for 3 d did not alter MØ density (Figure S8) but could efficiently increase the density of transepithelial protrusions derived from juxtatubular MØ in the medulla (Figure 7A) as well as their expression of Talin2 (Figure 7B). Next, we conducted proof-of-concept studies to investigate whether Piezo1 agonists could be used to treat kidney stones. We first adopted a prophylactic approach to investigate whether stimulating Piezo1 could attenuate renal stone formation (Figure 7C). To this end, wild-type adult mice were treated with an intravesical infusion of Yoda1 one day ahead and during the 3-d NaOx regimen. As expected, the control mice developed CaOx stones on harvesting, as assessed by Alizarin Red S staining (Figure 7D). Interestingly, the cohort of mice with intravesical Piezo1 treatment had only ∼50% of stone-affected area in comparison to the controls. Consistent with the notion that stone-induced tissue damage is associated with kidney inflammation (36), mice under Yoda1 treatment had reduced pro-inflammatory cytokine production in the kidney (Figure 7E). The protective effect of Yoda1 on kidney stone development was due to an action on Piezo1 expressed by medulla MØ, since *Piezo1*^ΔMØ^ mice, when received the same Yoda1 therapy, had no beneficial outcome relative to their vehicle-infused littermates (Figure 7F). Thus, prophylactic approach could limit stone generation.

**Figure 7.**
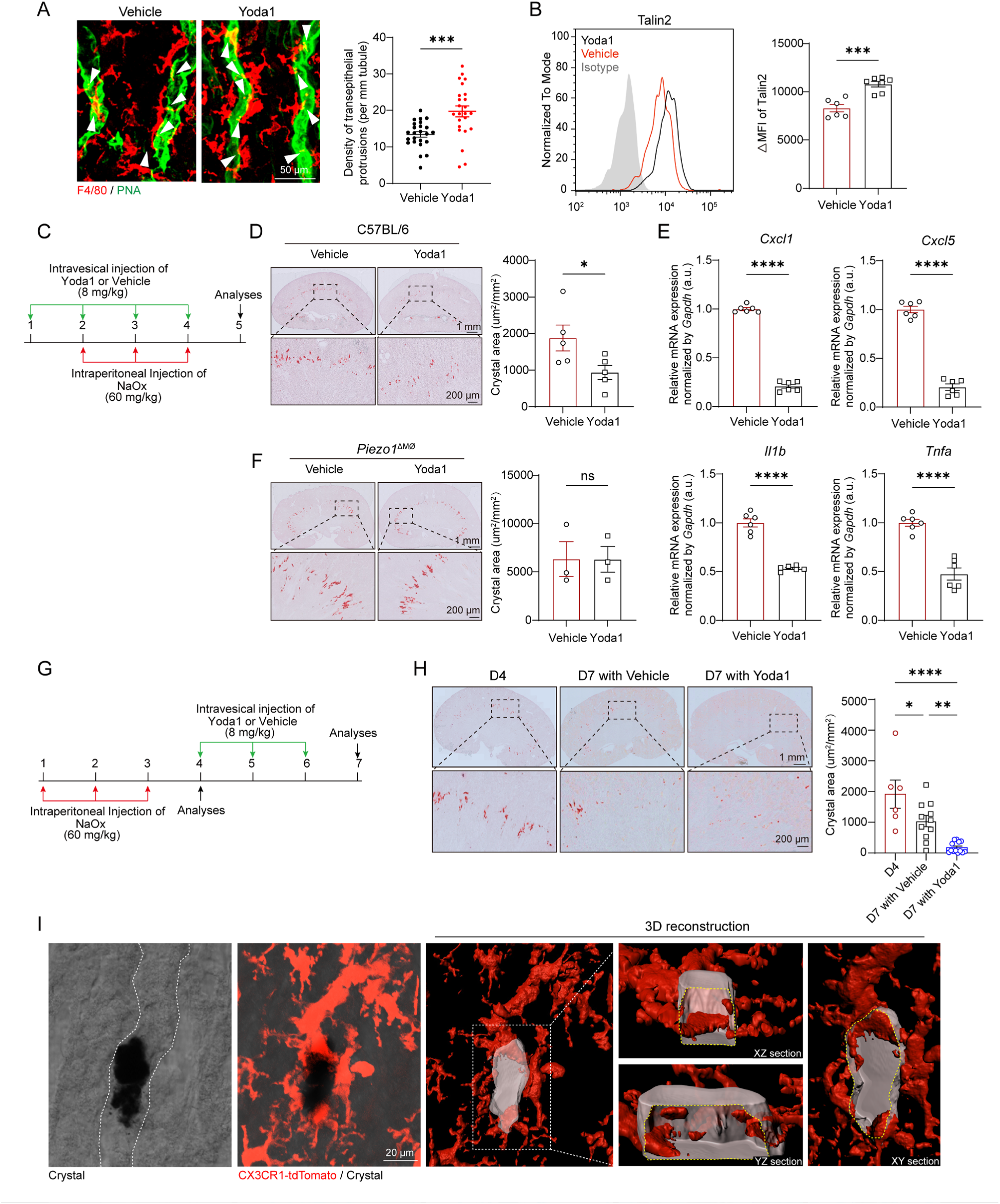
Stimulating macrophage Piezo1 with Yoda1 is valid to treat nephrolithiasis. (**A** and **B**) C57BL/6 mice received intravesical delivery of Yoda1 or vehicle once a day for 3 d. (**A**) The density of MØ-derived transepithelial protrusions on medullary collecting ducts was compared. Z-projections of 20 μm. Each dot represents the quantification from one 319 × 319 μm^2^ FOV. n = 6. (**B**) The expression of Talin2 in medulla MØ was compared by flow cytometry. (**C**) Scheme illustrating the prophylactic approach of experiments in Panels D-F. (**D**) In the oxalate nephrolithiasis model, some C57BL/6 mice received the prophylactic regimen of Yoda1 or vehicle as a control. Nephrolith-affected area were examined by Alizarin Red S staining and were compared. Each dot represents the average of the affected areas of two whole-mount slides from one mouse. (**E**) After the same procedure described in Panel D, the indicated pro-inflammatory chemokines and chemokines in the kidneys were examined by RT-PCR. (**F**) In the oxalate nephrolithiasis model, *Piezo1*^ΔMØ^ mice received the prophylactic regimen of Yoda1 or vehicle as a control. Nephrolith-affected area were examined and compared. Each dot represents the average of the affected areas of two whole-mount slides from one mouse. (**G**) Scheme illustrating the curative protocol of experiments for Panel H. (**H**) After the oxalate nephrolithiasis model, one group of C57BL/6 mice were sacrificed on the peak of the disease (Day 4) for nephrolith examination, and the other two groups of C57BL/6 mice were then received the curative regimen of Yoda1 or vehicle as a control. On Day 7, mice receiving the Yoda1 or vehicle were examined for nephrolith-affected area. Each dot represents the average of the affected areas of two whole-mount slides from one mouse. (**I**) MØ reporter *CX3CR1*^CreERT2/+^: Ai14 mice were treated with NaOx for 3 consecutive days to develop nephrolith. On Day 4, they received one intravesical infusion of Yoda1. Their kidneys were examined on Day 5 by Alizarin Red S staining. A representative image was shown. The brightfield image was obtained using the External Scanning Imaging Detector (ESID) channel of a confocal microscope. White dotted line, the contour of an affected kidney tubule. The magnifications show the corresponding 3D reconstructions viewed from different directions. Z-projections of 20 μm. n.s., not significant. \**P* < 0.05, \*\**P* < 0.01; \*\*\**P* < 0.001, \*\*\*\**P* < 0.0001 by two-tailed unpaired t test in (A, B, D, E, F) and one-way ANOVA in (H). Data are depicted as mean ± SEM. Data are derived from at least 2 independent experiments.

It is not that surprising that medulla MØ would remove miniature mineral crystals in the early stage of nephrolithiasis so that the above prophylactic approach could succeed. Considering that most patients with nephrolithiasis are not diagnosed until kidney stones have been large enough to be detected by imageology, e.g., ultrasound, or have caused symptoms, we wonder how efficient the Piezo1-targeting cell-based therapy would perform in treating established large stones. To answer this question, we next conducted a recovery protocol by delivering Yoda1 after kidney stone model was established. To this end, NaOx was given to 3 cohorts of mice for 3 consecutive days to let the mice develop CaOx stones. One group was examined one day after the last treatment (day 4) to evaluate the climax of kidney stones. The other two groups were followed with a 3-d regimen of Yoda1 or vehicle treatment (Figure 7G). It exhibited that mice would gradually recover from the disease since stone-affected area shrank to 54.4% in the vehicle-treated control mice on euthanasia (day 7) in comparison to the peak of the lesion (day 4) (Figure 7H). Remarkably, Yoda1 treatment greatly accelerated the recovery from nephrolithiasis in that the affected area was only 9.3% of the control on euthanasia (Figure 7H). To understand how medulla MØ coped with the established large mineral stones, the kidneys were collected from the mice receiving one dose of Yoda1 after the 3-d NaOx treatment (Figure 7I). We employed confocal analysis of thicker sections (30 μm) with high-resolution and optical magnification, and observed that the intratubular stones were generally surrounded by multiple MØ. Three-dimensional (3D) reconstruction revealed that MØ actually inserted protrusion into the stone, dissolving and removing the crystals bit by bit, instead of forming phagocytosing cups to intake the stone as a whole (Figure 7I). Altogether, our study for the first time indicates that medulla MØ-based therapy has a great potential to be developed for treating kidney stone diseases.

## DISCUSSION

Treatment options for patients with kidney stones include surgical stone removal and medication. Instrument miniaturization has greatly improved the surgical treatment of kidney stones (37). By contrast, medical management has not substantially advanced over the past 30 years and relies primarily on medications that modify urinary chemistry to reduce the supersaturation of crystallizing salts (4, 38). The physiological mechanisms that prevent kidney stone formation must be clarified to enable the development of novel targeted therapies.

We recently uncovered that kidney medulla MØ routinely surveyed the tubules and removed sedimentary particles in kidney tubules via their transepithelial protrusions (5). This finding provides an unprecedented opportunity to develop cell-based therapeutics for nephrolithiasis, which nevertheless requires in-depth understanding of the molecular mechanisms underlying MØ behavior and protrusion formation. We initiated the current study for this purpose and identified that mechanosensing via Piezo1 was an integral signal for medulla MØ to sense mechanical tension increase caused by tubule blockade, by mineral stones for instance. As a response, medulla MØ enhanced its tubule-cleaning ability by promoting transepithelial protrusion formation and possibly phagocytosis as well. Employing this finding, urinary tract delivery of an agonist of Piezo1 could stimulate the polarity of protrusion extension towards the tubules and facilitated their tubule “cleaning” functions. Thus, we successfully demonstrated here a proof-of-concept therapeutic strategy of treating nephrolithiasis by mobilizing the “cleaning” function of resident MØ in both prophylactic and curative approaches. In particular, thanks to the specificity of kidney targeting via urinary tract delivery (35), this method limited off-target side effects of Piezo1 stimulants on extrarenal organs to the maximum extent. However, this is just a start to understand the complicated mechanisms of grow transepithelial protrusions by kidney medulla MØ. As elaborate molecular pathways are expected to be elucidated in the near future, a wider range of MØ -associated molecular targets are set to be harnessed for treating nephrolithiasis.

Recently, mechanotransduction, or the process in which cells can convert physical forces into biochemical and electrochemical signals, has gained an increasing appreciation for its critical importance in regulating immunological processes (39). Piezo1 activation has been shown to promote efferocytosis (8, 17, 18) and enhance inflammation (19–22) in MØ of non-renal origin. However, our transcriptomic data of kidney medulla MØ was not in alignment with these paradigms, suggesting tissue heterogeneity of MØ biology. Instead, cell-matrix interaction, cell migration and cytoskeleton organization were among the highly enriched gene terms when Piezo1 was lost in medulla MØ, supporting the morphological alteration of MØ. In particular, we highlighted here that the expression of key players of integrin-dependent cytoskeleton remodeling, such as *Tln2* and *Ptk2*, were impaired when Piezo1 was absent, at least partially explaining the impairment of protrusion formation in Piezo1-deficient MØ. To control the structure, shape and behavior of cells in an ECM-directed manner, integrins form a functional biomechanical unit with the intracellular actin cytoskeleton (40). The cytoplasmic tail mediates its structural and signaling functions by binding to intracellular adaptor proteins, such as Talins, which assemble highly dynamic macromolecular focal adhesions (41). We show here that Piezo1 activation could upregulate Talin2, promote integrin β1 to take an activation conformation, and increase focal adhesion formation, which eventually mobilized cellular skeleton and enhanced protrusion formation. Talin family consists of two members, Talin1 and Talin2, which share 76% sequence identity (42). Talin1 is ubiquitously expressed and has been extensively studied (43, 44). In contrast, Talin2 is tissue-specific and has been shown to be expressed in heart, brain and skeletal muscle; the non-overlapping physiological functions of talin-2 relative to talin-1 remain largely uncharacterized (45), except that Talin1 is expressed at smaller focal adhesions in the peripheral region, whereas Talin2 is mainly found at large focal adhesions and fibrillar adhesions of fibroblasts (46–48). Here, we show that in medulla MØ, Talin2 but not Talin1 was inducible by Piezo1 in response to mechanical tension increase. Talin2 was mainly distributed at centrally located large focal adhesions in MØ and in the extended protrusions as well. Thus, Talin2 appeared not redundant to Talin 1 in MØ biology and mechanistically how it facilitates transepithelial protrusion formation warrants further investigation. To be noted, the promoted transepithelial protrusions post Piezo1 activation are much larger in size and length than the traditional plasma membrane protrusions such as lamellipodia, filopodia and blebs (49, 50), and are also different from podosome used by MØ for migration (28). In-depth study is required to elucidate the molecular details of their construction in the future.

## STUDY APPROVAL

All animal experiments adhered to the NIH Guide for the Care and Use of Laboratory Animals, and were approved by the Institutional Animal Care and Use Committee at Zhejiang University (ZJU20240259).

## ACKNOWLEDGMENTS

We thank Qin Han, Sanhua Fang, Shuangshuang Liu, Wei Yin, Qiong Huang, and Jingyao Chen from the Core Facilities, Zhejiang University School of Medicine, for the technical support. We also thank Zhengqi Liu from the Hangzhou Institute of Medicine (HIM), Chinese Academy of Sciences, for his assistance with the atomic force microscopy. This work was supported by grants from the Natural Science Foundation of Zhejiang Province (LZ26C110001 to X.Z.S., LY24H250003 to X.L., LQN26C080001 to J.H.), the National Natural Science Foundation of China (32470946 to X.Z.S., 82371252 to X.L.).

## AUTHOR CONTRIBUTIONS

Conceptualization, X.Z.S.; designing, R.H., X.L. and X.Z.S.; supervision, X.L. and X.Z.S.; imaging, R.H., Z.H., Y.L., N.C., Y.W. and J.H.; RNA-seq and bioinformatics analyses, G.C. and R.H.; blood biochemical analysis, Q.W.; all other experiments, R.H., Z.H., Y.L., and X.W.; statistics, R.H., Z.H., Y.L.; validation X.L.; writing, X.Z.S.

## METHODS

### Sex as a biological variable

Our study examined male and female animals, and similar findings are reported for both sexes.

### Mice

*Cx3cr1*^GFP/+^ [005582], *Cx3cr1*^CreERT2^ [020940], *Rosa26-stop-TdTomato* (Ai14) [007914], RCL- GCaMP6f (Ai95) [024105] and *Piezo1*^fl/fl^ [029213] mice were from The Jackson Laboratory. All the mice are in C57BL/6 background. C57BL/6 mice were purchased from Beijing Vital River Laboratory Animal Technology Co., Ltd. All mice used in this study without specific explanation were 8- to 12-week-old males. Mice receiving intravesical infusion were females. Mice were housed in a standard animal facility, with a 12-hr light/dark cycle, in specific-pathogen-free environment. All animal experiments adhered to the NIH Guide for the Care and Use of Laboratory Animals, and were approved by the Institutional Animal Care and Use Committee at Zhejiang University.

### In vivo treatments

To induce deletion of targeted genes, adult mice received intraperitoneal (i.p.) injection of 75 mg/kg (body weight) tamoxifen (MedChemExpress, #HY-13757A) daily for 7 consecutive days or regimens specified in experimental protocols.

Yoda1 (MedChemExpress, # HY-18723), a selective Piezo1 agonist, was first dissolved in DMSO and then diluted by 10 volumes in saline, which contained 20% of SBE-β-CD (absin, # abs816067) by weight. A dose of 8 mg/kg per day was administered

### Intrapelvic injection of latex beads

Mice were anesthetized and kept on a homeothermic pad to maintain body temperature during the procedure. Dorsal incision was made to expose kidney. The adipose tissue surrounding the kidney was gently separated to expose the pelvis which locates in the renal hilus. For intrapelvic injection, 25 μl 0.25% fluorescent latex beads (0.5 μm in diameter) (Sigma-Aldrich, #L3280) in Normal Saline was injected to the pelvis by an insulin syringe with a 30-gauge needle. The needle was kept in place for 10 seconds to forestall backflow. After that, the needle was slowly pulled back. The kidney was then returned to the abdominal cavity.

### Intravesical infusion

Intravesical infusion was performed on female mice, as described previously (35). Briefly, mice were water-deprived for 4 h prior to the procedure to minimize urine accumulation. Anesthesia was induced by i.p. injection of pentobarbital Sodium (80 mg/kg), and body temperature was maintained using a homeothermic heating pad. A catheter was gently inserted into the bladder via the urethra, and ∼60 μl vehicle or Yoda1 solution was slowly infused over approximately 30 s. The catheter was left in place for an additional 15 s to prevent backflow. After infusion, mice were placed in a supine position with hind limbs elevated at 60° until recovery from anesthesia (approximately 1–2 h).

### Immunohistological staining

For tissue immunofluorescence, mice were deeply anesthetized with isoflurane and transcardially perfused sequentially with ice-cold phosphate-buffered saline (PBS) and 4% paraformaldehyde (PFA), 5 min each. Kidneys were removed, post-fixed in 4% PFA overnight at 4 °C, and then dehydrated in PBS containing 30% (w/v) sucrose at 4 °C for 24 h. Frozen sections (30-µm thickness) were cut on a cryostat (CM3050S; Leica). Sections were blocked with 5% donkey serum in PBS for 2 h at room temperature (RT), followed by incubation with primary antibodies in antibody dilution buffer (0.25% Triton X-100 and 1% BSA in PBS) overnight at 4 °C, and then with appropriate secondary antibodies in the same buffer for 2 h at RT in the dark.

### Immunofluorescent staining of cultured cells

Cells grown on coverslips were fixed with 4% PFA for 10 min at RT, permeabilized with 0.1% Triton X-100 for 10 min, and blocked with 5% BSA in PBS for 30 min, all at RT. Cells were then incubated with primary antibodies overnight at 4 °C, followed by secondary antibodies for 2 h at RT in the dark.

All stained sections and coverslips were mounted with Fluoromount-G anti-fade mounting medium containing DAPI (Southern Biotech, #0100-20) and imaged using a confocal microscope (LSM 900; Carl Zeiss). Images were analyzed using Imaris (Bitplane) and Fiji (NIH) software.

The following antibodies were used: Rabbit anti-SLC12A/NKCC2 (Abcam, #ab191315); Rabbit anti-AQP1 (Millipore, #AB2219); Rabbit anti-AQP2 (Abcam, #ab199975); Rat anti-F4/80 (BM8) (ThermoFisher Scientific, #14 4801-81); Goat anti-KIM-1 (R&D Systems, #AF1817); Peanut Agglutinin (PNA)-CY5 (VectorLaboratories, #CL-1075-1); SF488 Phalloidin (Solarbio, #CA1640); Rabbit anti-Vinculin (HuaBio, #ET1705-94); Mouse anti-

Talin2 (Abcam, #ab105458); Rat Anti-CD29 (9EG7) (BD Biosciences, #550531); Donkey anti-rabbit IgG AF488 (Abcam, #ab150073); Donkey anti-rabbit IgG AF568 (Abcam, #ab175470); Donkey anti-rat AF647 (Abcam, #ab150155).

### Tissue disintegration and flow cytometry analysis

To analyze renal immune cells, kidneys were harvested from mice following transcardial perfusion with ice-cold PBS and mechanically minced into small fragments. In some experiments, the renal cortex and medulla were separated under a dissecting microscope (Olympus SZ-61-60). Each kidney sample was digested in 6 mL of RPMI 1640 medium containing 1.5 mg/mL collagenase IV (Worthington, #LS004189) and 100 U/mL DNase I (Sigma-Aldrich, #DN25) at 37 °C for 30 min with gentle agitation. After digestion, the cell suspension was filtered through a 70-µm strainer and subjected to density gradient centrifugation using 72% and 36% Percoll (Cytiva, #17089109). Immune cells were enriched at the interface between the two Percoll layers and subsequently collected. The isolated cells were analyzed by flow cytometry on a three-laser system (Agilent NovoCyte). Data were processed with FlowJo (v10.0).

The following antibodies were used for flow cytometry analysis: APC anti-mouse/human CD11b (M1/70) (Biolegend, #101211), Pacific Blue anti mouse/human CD11b (M1/70) (Biolegend, #101223), PE anti-mouse/human CD11b (M1/70) (Biolegend, #101207), BV510 anti-mouse CD45 (30-F11) (Biolegend, #103137), FITC anti mouse CD45 (30-F11) (Biolegend, #103107), APC/Fire750 anti-mouse F4/80 (BM8) (Biolegend, #123152), PE/Cyanine7 anti-mouse F4/80 (BM8) (Biolegend, #123113), Pacific Blue anti-mouse Ly-6C (HK1.4) (Biolegend, #128013), APC/Cyanine7 anti-mouse Ly-6G (1A8) (Biolegend, #127623), Brilliant Violet 605™ anti-mouse CD24 (Biolegend, #101827); APC anti-mouse/rat Integrin β1 (HMβ1-1) (Biolegend, #102215); Rabbit anti-Talin2 (ABclonal, # A19810); Rat Anti-CD29 (9EG7) (BD Biosciences, #550531); Rabbit anti- PIEZO1 (Novus Biologicals, # NBP1-78446); Donkey anti-rabbit AF488 (Abcam, #ab150073); Donkey anti rabbit AF647 (Abcam, #ab150075);

### Purification of renal macrophages for ex vivo assays

Kidney immune cells were first enriched by Percoll gradient centrifugation as described above and subsequently resuspended in MACS buffer (D-PBS supplemented with 2% FBS and 1 mM EDTA). After incubation with an Fc receptor blocking reagent (BioLegend, #156603), cells were stained with APC-conjugated anti-mouse F4/80 (BM8) antibody (BioLegend, #123115) on ice for 15 minutes, followed by washing with MACS buffer. Next, anti-APC Nanobeads (BioLegend, #488072) were added, and the cells were incubated on ice for an additional 15 minutes. Macrophages were then positively selected using a magnetic separation system and washed with PBS prior to downstream applications.

### RNA isolation and quantitative RT‒PCR

Tissue/cells were lysed using TRIzol reagent (Invitrogen), followed by chloroform. After phase separation, the samples were centrifuged at 12,000 × g for 15 min at 4 °C. The aqueous phase was carefully transferred into RNase-free tubes using RNase-free pipette tips. RNA was precipitated by adding 500 μl isopropanol. The resulting RNA pellet was washed with 75% ethanol and then resuspended in RNase-free water.

Total RNA from tissues was extracted using an ES Science kit (#ES-RN002plus), while RNA from cells was isolated using another ES Science kit (#ES-RN001plus). RNA concentrations were measured with a NanoDrop spectrophotometer. Subsequently, RNA was reverse-transcribed into cDNA using a high-capacity cDNA reverse transcription kit (Yeasen, #11141ES60). Quantitative real-time PCR (RT-PCR) was performed using SYBR Green Mix (Yeasen, #11201ES08) to assess mRNA levels of target genes. Gene expression levels were normalized to the housekeeping gene *Gapdh* and calculated using the 2^−ΔΔCt^ method. The following primers were used:

*Gapdh*: Forward primer AGGTCGGTGTGAACGGATTTG

Reverse primer TGTAGACCATGTAGTTGAGGTCA

*Piezo1*: Forward primer AGGACTTCCCCACCTATTGG

Reverse primer CCAGGGATGAGGATACTGGAAAA

*Piezo2*: Forward primer ATGTGCGTTCCGGTACAATGG

Reverse primer TGTGTCCTTGCATCGTTGCT

*Kcnk2*: Forward primer CCGAGGCTCTCATTCTCCTCA

Reverse primer AGGACGACCACCAGGAAAATC

*Kcnk4*: Forward primer ATCTGGGGCTCTAGTGTTCCA

Reverse primer CCAAGCTGATGAGTGGTTGCT

*Kcnk10*: Forward primer TGGCTGCATCGTGTTTGTGA

Reverse primer CTGTGGTCAGCGTGACTACC

*Kcnk18*: Forward primer CTCTCTTCTCCGCTGTCGAG

Reverse primer AAGAGAGCGCTCAGGAAGG

*Trpa1*: Forward primer GTCCAGGGCGTTGTCTATCG

Reverse primer CGTGATGCAGAGGACAGAGAT

*Trpv4*: Forward primer ATGGCAGATCCTGGTGATGG

Reverse primer GGAACTTCATACGCAGGTTTGG

*Stoml3*: Forward primer GATTCACCGGAGAAACTGGAG

Reverse primer TCCATACTGAGATTGGGAAGGT

*Itgb1*: Forward primer ATGCCAAATCTTGCGGAGAAT

Reverse primer TTTGCTGCGATTGGTGACATT

*Col1a1*: Forward primer CTGACGCATGGCCAAGAAGA

Reverse primer CGTGCCATTGTGGCAGATAC

*Col3a1*: Forward primer CTGTAACATGGAAACTGGGGAAA

Reverse primer CCATAGCTGAACTGAAAACCACC

*Fn1*: Forward primer TTCAAGTGTGATCCCCATGAAG

Reverse primer CAGGTCTACGGCAGTTGTCA

*Tln2*: Forward primer GGACCTGGGAAGCACCTCTAA

Reverse primer TGTGTAATGCTCATTGCCTTGA

*Ptk2*: Forward primer GAGTACGTCCCTATGGTGAAGG

Reverse primer CTCGATCTCTCGATGAGTGCT

*Slc34a1*: Forward primer TGCCTCTGATGCTGGCTTTC

Reverse primer GATAGGATGGCATTGTCCTTGAA

*Havcr1*: Forward primer GTTAAACCAGAGATTCCCACACG

Reverse primer TCTCATGGGGACAAAATGTAGTG

*Cxcl1*: Forward primer CTGGGATTCACCTCAAGAACATC

Reverse primer CAGGGTCAAGGCAAGCCTC

*Cxcl5*: Forward primer GGTCCACAGTGCCCTACG

Reverse primer GCGAGTGCATTCCGCTTA

*Il1b*: Forward primer ACCTTCCAGGATGAGGACATGA

Reverse primer CTAATGGGAACGTCACACACCA

*Tnfa*: Forward primer AAGCCTGTAGCCCACGTCGTA

Reverse primer GGCACCACTAGTTGGTTGTCTTTG

### Ex vivo calcium Imaging and the mechanical brushing assay

Primary medulla macrophages isolated from *CX3CR1*^CreERT2/+^: GCaMP6f mice or *CX3CR1*^CreERT2/+^: GCaMP6f: *Piezo1*^fl/fl^ mice were seeded onto 35-mm confocal dishes (NEST). Cells were incubated in HBSS containing Ca²⁺ and Mg²⁺ (Solarbio) for 30 min at 37°C in a dark incubator for environmental adaptation. Real-time calcium imaging was performed using a confocal microscope (LSM 900; Carl Zeiss). To record Yoda1-induced calcium influx, cells were stimulated with 10 µM Yoda1. For mechanical stimulation, a 0.008-g von Frey filament (Bioseb) was brushed across the cell surface. Fluorescence intensity changes (GCaMP6f) were recorded in real time and analyzed to reflect intracellular calcium dynamics.

### Polyacrylamide hydrogel preparation

The preparation of polyacrylamide (PA) hydrogel with varying stiffness was adapted from previous studies (51–54). Round glass coverslips (NEST) were designated as bottom or top coverslips. Bottom coverslips were coated with 1.2% bind silane (Yeasen, #36241ES25) dissolved in 95% ethanol and 5% glacial acetic acid, and air-dried. Top coverslips were incubated in 15% Sigmacote (Sigma-Aldrich) dissolved in chloroform for 1 h, air-dried, and polished.

PA hydrogel with different stiffness were prepared by mixing different concentrations of acrylamide (Acr) and bis-acrylamide (Bis-Acr). For soft gels (1–4 kPa), a solution of 8% Acr and 0.05% Bis-Acr (Sigma-Aldrich) was prepared in ddH₂O. For stiff gels (62–68 kPa), 8% Acr and 0.7% Bis-Acr were used. Polymerization was initiated by adding TEMED and ammonium persulfate. The mixed acrylamide solution was rapidly pipetted onto the bottom coverslip, and the top coverslip was placed on the semi-solidified gel. After polymerization, the top coverslip was removed, leaving the gel adhered to the bottom coverslip. Gels were then immersed in 10 mM HEPES buffer containing 2% (v/v) penicillin/streptomycin for at least 48 h.

For surface functionalization, gels were treated with 0.2 mg/mL Sulfo-SANPAH (Sigma-Aldrich) and exposed to long-wave UV light (365 nm) for 10–25 min. After UV activation, gels were washed with 50 mM HEPES buffer and incubated with 0.2 mg/mL collagen type I and 0.1 mg/mL fibronectin overnight at 4°C. The functionalized gels were washed three times with PBS before cell seeding. All steps were performed under aseptic conditions on a UV-sterilized clean bench.

### Biological atomic force microscope

In this study, the Young’s modulus of kidney tissue was measured using Biological Atomic Force Microscope (BioAFM). BioAFM indentation was performed with a NanoWizard 4 XP (Bruker) in Quantitative Imaging (QI) mode using PFQNM-LC-V2 probes. Cantilever spring constants and deflection sensitivity were calibrated prior to each measurement session. Force-distance curves were acquired in liquid environment to maintain tissue hydration. The apparent Young’s modulus was extracted by fitting the force-indentation curves with the Hertz model (Sneddon modification), assuming a Poisson’s ratio of 0.5. All modulus fitting calculations were performed using JPKSPM Data Processing software (Bruker).

### Live-Cell imaging of macrophage dynamic protrusion formation

Primary medulla macrophages isolated from *Cx3cr1*^GFP/+^ mice were pretreated with 1 µM Yoda1 or DMSO (vehicle control) for 24 h. Afterwards, cells were replated onto 35-mm confocal dishes coated with 0.1 mg/mL fibronectin. Following a 30-min attachment period, real-time fluorescence imaging was performed using a confocal microscope (LSM 900; Carl Zeiss) equipped with a live-cell incubation system. GFP fluorescence was recorded continuously for 1 h to monitor dynamic protrusion changes and cell morphology.

### Calcium oxalate kidney stone model

Mice received i.p. injections of sodium oxalate (NaOx) (Sigma-Aldrich, #71800) at a dose of 60 mg/kg once daily for three consecutive days, or according to regimens specified in individual experimental protocols. Given that female mice are generally smaller than males and exhibit reduced tolerance to drug administration, we used 12- to 16-week-old females for all experiments.

### Analyses of mineral crystals in the kidney

von Kossa staining and Alizarin Red S staining was used to detect calcium deposits. For von Kossa staining, mice were anesthetized and transcardially perfused with ice-cold PBS for 5 min, followed by 4% PFA for an additional 5 min. Kidneys were then harvested and post-fixed in 4% PFA for 48 h. Paraffin-embedded sections were prepared and subjected to staining. For visualization of large calcium oxalate (CaOx) stones, frozen sections were incubated with 0.2% Alizarin Red S solution (pH 6.5–6.75) at 37 °C for 10 min.

To examine CaPi microcrystals, mice received an intravenous injection of OsteoSense 680EX (Revvity, #NEV10020EX) at 48 nmol/kg 30 min prior to transcardial perfusion. Subsequently, tissue slices of 60-μm thickness were prepared for imaging analysis.

### Blood and urine biochemical analyses

Blood samples were collected via retro-orbital bleeding, and plasma was separated by centrifugation at 1,500 × g for 15 min at 4 °C. Urine samples were obtained from mice housed individually in metabolic cages over a 24-h collection period. Samples were kept on ice throughout collection, and total urine volume was recorded. Urine samples were then centrifuged at 1,000 × g for 20 min at 4 °C, and the supernatant was collected for subsequent analysis.

Concentrations of phosphorus (Pi) and calcium (Ca²⁺) in both plasma and urine were measured using Cobas 8000 ISE and Cobas c702 analyzer.

### Bone marrow-derived macrophages (BMDMs) culture

Bone marrows were isolated from the femurs and tibias of 6- to 8-week-old C57BL/6 mice, an then cultured in RPMI 1640 medium supplemented with 10% fetal bovine serum (FBS), 1% penicillin-streptomycin, 2 mM L-glutamine (Solarbio, #G0200), 10 mM HEPES (Sangon, #E60701), and 15% L929 cell-conditioned medium as a source of macrophage colony-stimulating factor (M-CSF). The cells were maintained under standard culture conditions, with fresh medium replaced on day 3 and day 6. On day 7, the differentiated BMDMs were harvested, seeded into culture plates, and used for subsequent experiments.

### Liquid Chromatography-Mass Spectrometry (LC-MS)

Blood samples were collected via cardiac puncture prior to perfusion, and tissue samples were harvested after perfusion with ice-cold Normal Saline. For plasma preparation, blood was collected in EDTA-containing tubes and centrifuged at 1,500 × g for 20 min at 4°C to separate plasma. For tissue samples, an appropriate amount of kidney tissue was homogenized in 50% aqueous methanol at a ratio of 4:1 (mL/g tissue) using a tissue homogenizer to obtain tissue homogenates.

For LC-MS/MS analysis, 50 µL of each sample (plasma or tissue homogenates) was mixed with 400 µL of acetonitrile containing 2 ng/mL verapamil (internal standard). The mixture was vortexed at 2,500 rpm for 3 min and centrifuged at 15,000 × g for 10 min at 4°C. The supernatant was collected for LC-MS/MS analysis.

Chromatographic separation was performed on an ACQUITY UPLC® BEH C18 column (2.1 × 50 mm, 1.7 µm; Waters) using a mobile phase consisting of water with 0.1% formic acid (A) and acetonitrile (B). The LC system was coupled to an API4000 mass spectrometer (AB SCIEX QTRAP 5500) equipped with an electrospray ionization (ESI) source operating in positive ion mode. Data acquisition and analysis were carried out using Analyst 1.6.3 software (AB Sciex, Ontario, Canada).

### RNA-seq library construction

Medulla macrophages were purified by cell sorting with a BD SORP ARIA II. Holo-Seq, a recently developed method designed for improving data fidelity of low-cell-number samples (55). Following the Holo-Seq method, RNA-seq libraries were constructed from renal macrophages. Briefly, the collected cells were then lysed in RLT Plus buffer supplemented with 1% β-mercaptoethanol and 100 ng of a synthetic RNA spike-in mixture. mRNA was subsequently enriched using the NEBNext Poly(A) mRNA Magnetic Isolation Module. Afterward, Not I restriction enzyme was added to digest the synthetic carrier RNA, and cDNA was synthesized. Sequencing libraries were then prepared according to the protocol of the Ultra RNA Library Prep Kit for Illumina. The resulting DNA was amplified and purified by PCR, and its quality was assessed using an Agilent 2200 Bioanalyzer. Finally, libraries that passed quality control were sequenced on the Illumina HiSeq X Ten platform.

### RNA-seq data analysis

RNA-seq data analysis was performed as follows. Adapter and quality trimming of raw sequencing reads were carried out using BBDuk (BBMap v38.86), removing regions with an average quality score below 15 and discarding reads shorter than 36 bp. Cleaned reads were then aligned to the mouse genome (GRCm38, GENCODE annotation) using STAR (v2.7.5a_2020-6-19) with default parameters in paired-end mode. Transcript abundance was quantified and normalized using the ‘quant’ module of Salmon (v1.2.1). Differential expression analysis was subsequently performed using the R package DESeq2 (v1.34.0) to compare medullary macrophages isolated from *Piezo1*^ΔMØ^ mice versus those from control mice. This analysis yielded 280 significantly upregulated genes (log_2_FC > 1, P < 0.05) and 1,083 downregulated genes (log_2_FC < –1, P < 0.05). Finally, Gene Ontology (GO) enrichment analysis of the differentially expressed genes was performed using the clusterProfiler package (v4.0.5) in R.

### Statistics

Statistical analysis was performed with Prism 8.0 (GraphPad). Data are presented as mean ± SEM. One-way ANOVA with Tukey’s multiple-comparisons testing was used to compare multiple groups. Unpaired Student’s t tests were used to compare two groups. All statistical tests were two tailed, and *P* values of <0.05 were considered significant.

