## Supplemental Figures for "Kidney medulla macrophages maintain a free flow of urine by sensing force"

Figure S1

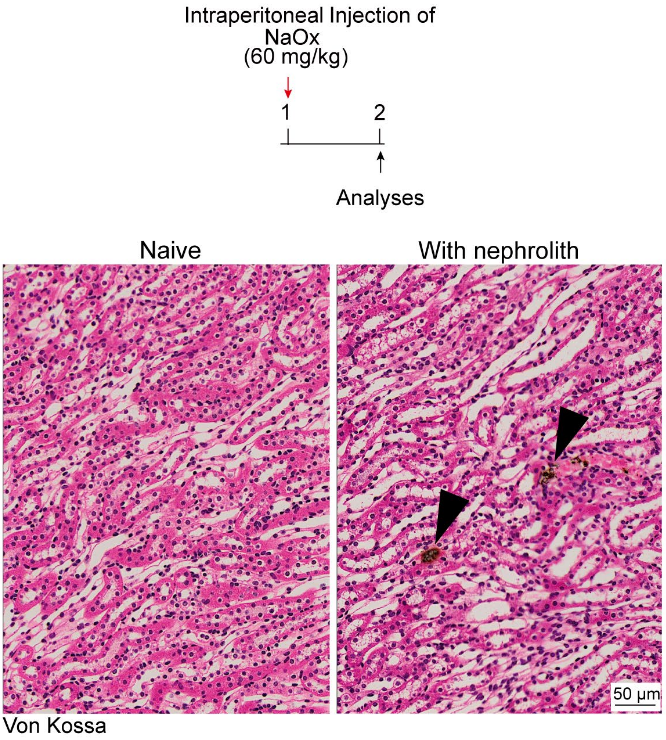


**Figure S1. The oxalate nephrolith model.** C57BL/6 mice were i.p. treated with 60 mg/kg NaOx and the kidneys were examined 24 hr later by Von Kossa staining. The representative images of normal kidney and the kidney with nephrolith were shown. Arrowhead, CaOx crystals.

Figure S2

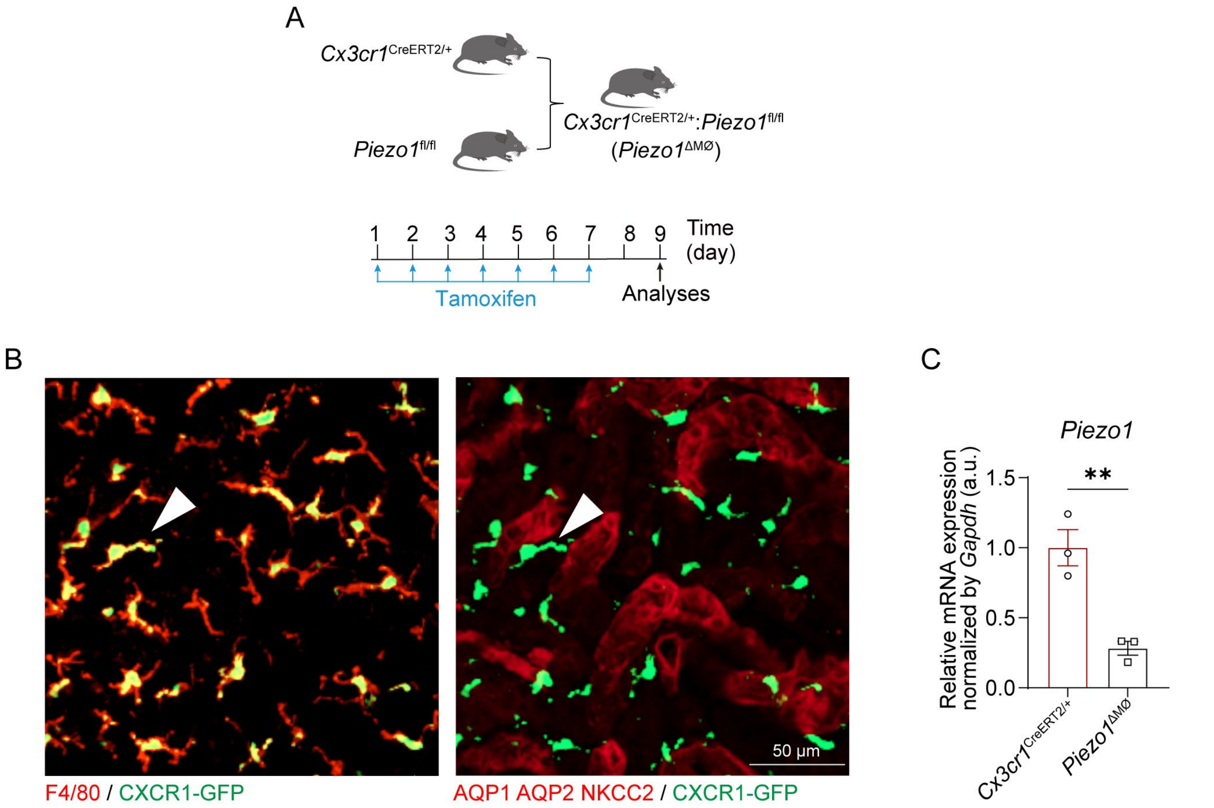


**Figure S2. Generation and validation of *Piezo1*$^{\Delta MØ}$ mice**. (**A**) Strategy of generating *Piezo1*$^{\Delta MØ}$ mice. (**B**) *Cx3cr1*$^{GFP/+}$ mice show that GFP colocalized with F4/80$^{+}$ MØ but not tubular epithelial cells. AQP1, AQP2 and NKCC2 together label most medullary tubules. (**C**) Efficiency of *Piezo1* deletion in kidney medulla MØ, examined by RT-PCR. Each dot represents a pool of cells derived from 4 mice. $^{**}P < 0.01$ by two-tailed unpaired t test. Data are depicted as mean ± SEM.

### Figure S3

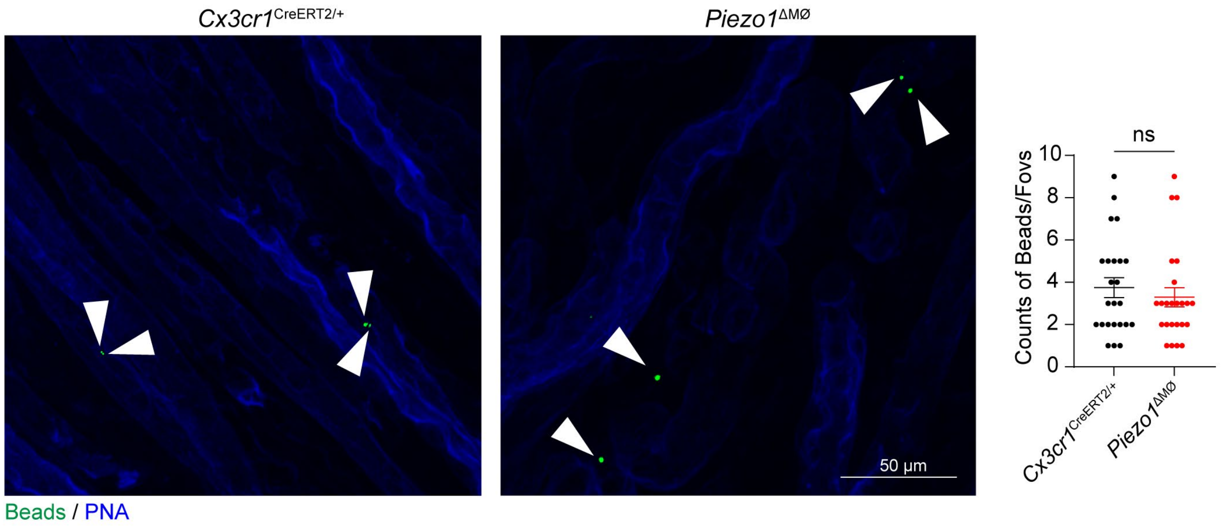


**Figure S3. Bead retention in the kidney medulla.** *Piezo1*$^{\Delta MØ}$ mice and their control mice received an intrapelvic injection of fluorescent beads. Eight hours later, bead densities in the medulla were counted and compared. Each dot represents the quantification from one 319 × 319 μm$^2$ FOV, n = 6. n.s., not significant by two-tailed unpaired t test. Data are depicted as mean ± SEM.

Figure S4

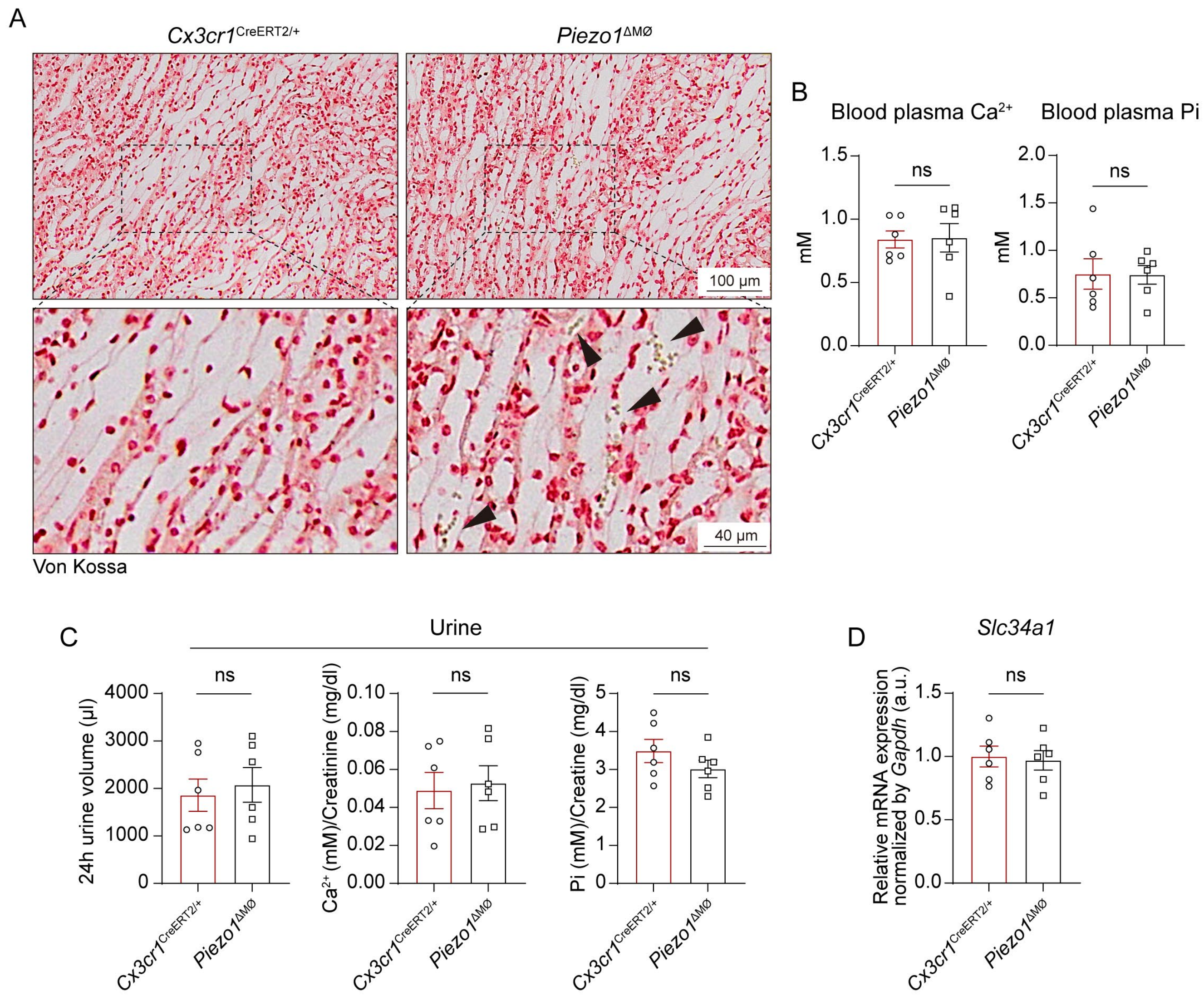


**Figure S4. The phenotypes of *Piezo1*$^{\Delta M\text{Ø}}$ mice**. *Piezo1*$^{\Delta M\text{Ø}}$ mice and the control mice were treated with tamoxifen as shown in Figure S2A. (**A**) Representative images showed mineral particle deposition in the medulla, examined by Von Kossa staining. Arrowhead, mineral particles. (**B**) Blood concentrations of calcium and phosphate. (**C**) 24-hr urine output, and the concentrations of calcium and phosphate in the urine, adjusted by the excreted creatinine. (**D**) Cortex tubule epithelial cells were purified and the expression of *Slc34a1* was examined by RT-PCR. n.s., not significant by two-tailed unpaired t test in. Data are depicted as mean ± SEM.

### Figure S5

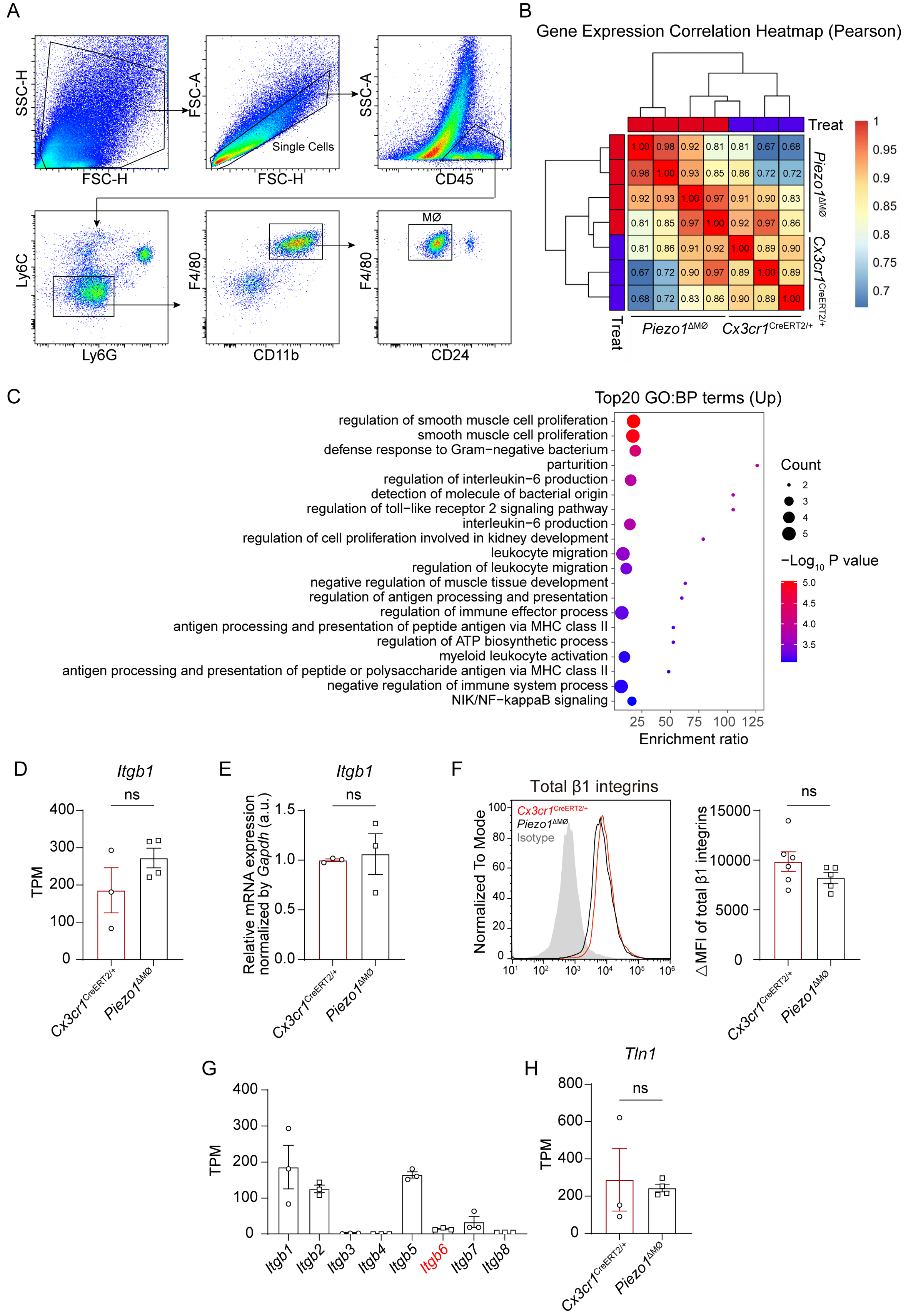

A
SSC-H
FSC-H
FSC-A
FSC-H
Single Cells
SSC-A
CD45
Ly6C
Ly6G
F4/80
CD11b
MØ
F4/80
CD24
B
Gene Expression Correlation Heatmap (Pearson)
Treat
Piezo1ΔMØ
Cx3cr1CreERT2/+
1.00 0.98 0.92 0.81 0.81 0.67 0.68
0.98 1.00 0.93 0.85 0.86 0.72 0.72
0.92 0.93 1.00 0.97 0.91 0.90 0.83
0.81 0.85 0.97 1.00 0.92 0.97 0.86
0.81 0.86 0.91 0.92 1.00 0.89 0.90
0.67 0.72 0.90 0.97 0.89 1.00 0.89
0.68 0.72 0.83 0.86 0.90 0.89 1.00
1 0.95 0.9 0.85 0.8 0.75 0.7
C
Top20 GO:BP terms (Up)
regulation of smooth muscle cell proliferation
smooth muscle cell proliferation
defense response to Gram−negative bacterium
parturition
regulation of interleukin−6 production
detection of molecule of bacterial origin
regulation of toll−like receptor 2 signaling pathway
interleukin−6 production
regulation of cell proliferation involved in kidney development
leukocyte migration
regulation of leukocyte migration
negative regulation of muscle tissue development
regulation of antigen processing and presentation
regulation of immune effector process
antigen processing and presentation of peptide antigen via MHC class II
regulation of ATP biosynthetic process
myeloid leukocyte activation
antigen processing and presentation of peptide or polysaccharide antigen via MHC class II
negative regulation of immune system process
NIK/NF−kappaB signaling
25 50 75 100 125
Enrichment ratio
Count
2 3 4 5
−Log10 P value
5.0 4.5 4.0 3.5
D
Itgb1
ns
TPM
0 100 200 300 400
Cx3cr1CreERT2/+
Piezo1ΔMØ
E
Itgb1
ns
Relative mRNA expression normalized by Gapdh (a.u.)
0.0 0.5 1.0 1.5
Cx3cr1CreERT2/+
Piezo1ΔMØ
F
Total β1 integrins
Cx3cr1CreERT2/+
Piezo1ΔMØ
Isotype
Normalized To Mode
0 20 40 60 80 100
10¹ 10² 10³ 10⁴ 10⁵ 10⁶
ns
ΔMFI of total β1 integrins
0 5000 10000 15000
Cx3cr1CreERT2/+
Piezo1ΔMØ
G
TPM
0 100 200 300 400
Itgb1 Itgb2 Itgb3 Itgb4 Itgb5 Itgb6 Itgb7 Itgb8
H
Tln1
ns
TPM
0 200 400 600 800
Cx3cr1CreERT2/+
Piezo1ΔMØ

**Figure S5. Transcriptomic analysis of medulla macrophages derived from *Piezo1*$^{\Delta MØ}$ mice**. (**A**) The gating strategy for kidney medulla MØ. (**B**) Correlation hierarchical clustering analysis of the experimental samples. (**C**) Top 20 upregulated pathways of *Piezo1-deprived* medulla MØ relative to control MØ, derived from GO analysis of DEGs. (**D**) *Itgb1* expression comparison between the medulla MØ derived from the indicated mice, according to gene expression values (TPM) of the transcriptomic data. (**E**) *Itgb1* expression in the indicated medulla MØ was evaluated by RT-PCR. (**F**) Total integrin β1 expression was evaluated by flow cytometry. (**G**) Relative expression levels of all known genes encoding integrin β subtypes according to TPM of medulla MØ derived from tamoxifen-treated *Cx3cr1*$^{CreERT2/+}$ mice. *Itgb6* is highlighted. (**H**) Relative expression levels of *Tln1* according to TPM of the transcriptomic data. Each dot represents a pool of cells derived from two mice in (D, G, H). Each dot represents a pool of cells derived from 4 mice in (E). n.s., not significant by two-tailed unpaired t test. Data are depicted as mean ± SEM.

Figure S6

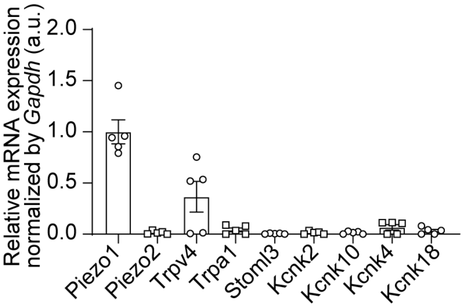


**Figure S6. Bone marrow-derived macrophages relatively highly express Piezo1.** The transcript expression profiles of mechanoreceptors in BMDM, assessed by RT-PCR. Each dot represents an individual mouse.

Figure S7

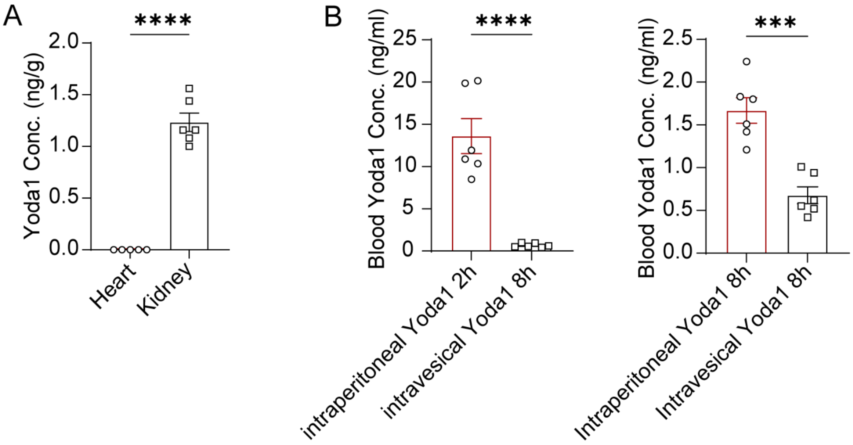


**Figure S7. The specificity of intravesical delivery of Yoda1 to the kidney.** (**A**) C57BL/6 mice received an intravesical injection of Yoda1 (8 mg/kg). Eight hours later, Yoda1 was examined by mass spectrometry in the heart and the kidney. (**B**) After the indicated routes of delivery, Yoda1 concentrations in the blood plasma were examined by mass spectrometry after the indicated time. ***$P < 0.001$, ****$P < 0.0001$ by two-tailed unpaired t test. Data are depicted as mean ± SEM.

Figure S8

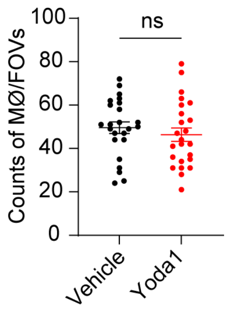


**Figure S8. Intravesical infusion of Yoda1 does not alter the density of kidney medulla macrophages.** C57BL/6 mice received intravesical delivery of Yoda1 or vehicle once a day for 3 d. On Day 4, kidney medulla were examined for F4/80 staining. Each dot represents the quantification of MØ from one 319 × 319 $\mu m^2$ FOV, n = 6. n.s., not significant by two-tailed unpaired t test. Data are depicted as mean ± SEM.
